# Distinct antibody repertoires against endemic human coronaviruses in children and adults

**DOI:** 10.1101/2020.06.21.163394

**Authors:** Taushif Khan, Mahbuba Rahman, Fatima Al Ali, Susie S. Y. Huang, Manar Ata, Qian Zhang, Paul Bastard, Zhiyong Liu, Emmanuelle Jouanguy, Vivien Béziat, Aurélie Cobat, Gheyath K. Nasrallah, Hadi Yassine, Maria Smatti, Amira Sayeed, Isabelle Vandernoot, Jean-Christophe Goffard, Guillaume Smits, Isabelle Migeotte, Filomeen Haerynck, Isabelle Meyts, Laurent Abel, Jean-Laurent Casanova, Mohammad R. Hasan, Nico Marr

**Affiliations:** Research Branch, Sidra Medicine, Doha, Qatar; St Giles Laboratory of Human Genetics of Infectious Diseases, Rockefeller Branch, The Rockefeller University, New York, NY, USA; Laboratory of Human Genetics of Infectious Diseases, Necker Branch, INSERM U1163, Necker Hospital for Sick Children, Paris, France, EU; University of Paris, Imagine Institute, Paris, France, EU; College of Health Sciences and Biomedical Research Center, Qatar University, Doha, Qatar; Department of Pathology, Sidra Medicine, Doha, Qatar; Center of Human Genetics, Hôpital Erasme, Université Libre de Bruxelles, Brussels, Belgium; Department of Internal Medicine, Hôpital Erasme, Université Libre de Bruxelles, Brussels, Belgium; Fonds de la Recherche Scientifique (FNRS) and Center of Human Genetics, Hôpital Erasme, Université Libre de Bruxelles, Brussels, Belgium; Department of Respiratory Diseases, Department of Pediatrics and Genetics, Center for Primary Immunodeficiencies Ghent, Jeffrey Modell Foundation diagnostic and research center, Gent, Belgium, EU; Laboratory for Inborn Errors of Immunity, Department of Microbiology, Immunology and Transplantation, Department of Pediatrics, University Hospitals Leuven, KU Leuven, Belgium; Department of Pediatrics, University Hospitals Leuven, KU Leuven, Belgium; Pediatric Hematology and Immunology Unit, Necker Hospital for Sick Children, AP-HP, Paris, France, EU; Howard Hughes Medical Institute, New York, NY, USA; Department of Pathology and Laboratory Medicine, Weill Cornell Medical College in Qatar, Doha, Qatar; College of Health and Life Sciences, Hamad Bin Khalifa University, Doha, Qatar

## Abstract

Four endemic human coronaviruses (HCoVs) are commonly associated with acute respiratory infection in humans. B cell responses to these “common cold” viruses remain incompletely understood. Here we report a comprehensive analysis of CoV-specific antibody repertoires in 231 children and 1168 adults using phage-immunoprecipitation sequencing. Seroprevalence of antibodies to endemic HCoVs ranged between ~4 and 27% depending on the species and cohort. We identified at least 136 novel linear B cell epitopes. Antibody repertoires against endemic HCoVs were qualitatively different between children and adults in that anti-HCoV IgG specificities more frequently found among children targeted functionally important and structurally conserved regions of the spike, nucleocapsid and matrix proteins. Moreover, antibody specificities targeting the highly conserved fusion peptide region and S2’ cleavage site of the spike protein were broadly cross-reactive with peptides of epidemic human and non-human coronaviruses. In contrast, an acidic tandem repeat in the N-terminal region of the Nsp3 subdomain of the HCoV-HKU1 polyprotein was the predominant target of antibody responses in adult donors. Our findings shed light on the dominant species-specific and pan-CoV target sites of human antibody responses to coronavirus infection, thereby providing important insights for the development of prophylactic or therapeutic monoclonal antibodies and vaccine design.

## Introduction

Four endemic human-tropic coronaviruses (HCoVs) are commonly associated with respiratory illness in humans, namely HCoV-229E, -NL63, -OC43, and -HKU1 [1–4]. Clinical outcomes of acute infection with these HCoVs range from mild upper respiratory tract infections in most patients, to viral bronchiolitis and pneumonia in rare patients, the latter requiring hospitalization [5]. The ratio of more severe versus mild outcomes of acute infection with endemic HCoVs is largely comparable to that of other “common cold” viruses, such as human respiratory syncytial virus (HRSV), human rhinoviruses (HRVs), human adenoviruses (HAdVs), and human parainfluenza viruses (HPIVs), albeit with differences in seasonality and prevalence of the viruses depending on the species [5–7]. In addition to the four endemic HCoV, three human-tropic epidemic coronaviruses (CoVs) have emerged over the last two decades, namely Severe Acute Respiratory Syndrome-associated CoV (SARS-CoV) [8], Middle East Respiratory Syndrome-associated coronavirus (MERS-CoV) [9], and SARS-CoV-2 [10], the etiological agent of Coronavirus Disease 2019 (COVID-19) which has now reached pandemic proportions [11]. Similar to endemic HCoVs, infection of humans with epidemic CoVs is associated with a wide range of outcomes but leads more frequently to severe clinical manifestations such as acute respiratory distress syndrome (ARDS) [12–14]. Phylogenetic analyses suggest that, similar to these epidemic CoVs, all endemic HCoVs are of zoonotic origin and their possible ancestors share similar natural animal reservoirs and intermediate hosts [6]. HCoV-229E may have been transferred from dromedary camels, similar to MERS-CoV, while HCoV-OC43 is thought to have emerged more recently from ancestors in domestic animals such as cattle or swine in the context of a pandemic at the end of the 19th century [6, 15].

The wide variability in transmissibility and clinical manifestations of infections by endemic and epidemic CoVs among humans remains poorly understood. On the population level, the case-fatality rate is highest for MERS (~34 - 37%) and several risk factors are associated with progression to ARDS in MERS, SARS and COVID-19 cases, including old age, diabetes mellitus, hypertension, cancer, renal and lung disease, and co-infections [12, 16]. Nonetheless, even MERS-CoV infection among humans can run a completely asymptomatic course in some cases, particularly among children [17–19]. There is evidence that children are generally less susceptible to infection with epidemic CoVs and once infected, they are less likely to experience severe outcomes compared to adults, although this important association and the underlying reasons remain poorly understood [12, 18, 20, 21]. Importantly, it remains unclear to what extent pre-existing immunity from past infections with endemic HCoVs provides some degree of cross-protection and affects clinical outcomes of infection with the epidemic SARS-CoV-2 or MERS-CoV. Our overall understanding of the immunity induced by natural infection with endemic HCoVs remains very limited. Serological studies have shown some degree of cross-reactive antibodies in patients with past CoV infections but many of these studies were limited in sample size and often focused on specific viral antigens only [22–25]. Depending on their binding affinity and specificities, such cross-reactive antibodies could either have no effect on clinical outcomes, may provide protection from severe disease to some degree, or may lead to antibody-dependent enhancement of disease—the latter can be a major obstacle in vaccine development [26]. Interestingly, two recent studies from independent groups have shown that a considerable proportion of individuals without a history of SARS-CoV-2 infection have SARS-CoV-2-reactive T cells, which suggests that cross-reactive T cell subsets originating from past infections by endemic HCoVs may play a role in the clinical course of infection with the phylogenetically related epidemic CoVs [27, 28]. A systematic assessment to elucidate the immunodominant B cell antigen determinants of endemic HCoVs has not been done. We hypothesized that a fraction of the general population also have antibodies generated during past encounters with ‘common cold’ coronaviruses that cross-react with proteins of epidemic CoVs. This may affect the dynamics of sporadic MERS outbreaks that mostly occur in the Middle East, and the current COVID-19 pandemic.

## Results

To gain a deeper insight into human antibody responses to endemic HCoVs, we performed phage-immunoprecipitation sequencing (PhIP-Seq) [29–31] on previously collected serum or plasma samples obtained from a total of 1431 human subjects from three different cohorts. These included i) healthy male adult blood donors (ABD) with diverse ethnic background and nationality (Supplementary Figure S1A); ii) adult male and female participants of a national cohort study—the Qatar Biobank (QBB) [32]—representing the general population (Supplementary Figure S1B); and iii) pediatric outpatients and inpatients who were tested for metabolic conditions unrelated to infection, chronic disease or cancer (Methods and Supplementary Figure S1C). The samples were collected prior to the current COVID-19 outbreak (Methods). In brief, PhIP-Seq allowed us to obtain comprehensive antiviral antibody repertoires across individuals in our three human cohorts using phage display of oligonucleotide-encoded peptidomes, followed by immunoprecipitation and massive parallel sequencing [29, 31]. The VirScan phage library used for PhIP-Seq in the present study comprised peptides derived from viral proteins—each represented by peptide tiles of up to 56 amino acids in length that overlap by 28 amino acids—which collectively encompass the proteomes of a large number of viral species, including HCoV-229E, -NL63, -HKU1 and -OC43 [29, 31]. Proteins of endemic HCoVs which were represented in the VirScan phage library included the Orf1ab replicase polyprotein (pp1ab), the spike glycoprotein (S), the matrix glycoprotein (M), the nucleocapsid protein (N) and gene products of the species- and strain-specific open reading frames (ORFs) encoded in the 3’ region of the viral genomes (Supplementary Table S1). Of note, we utilized an expanded version of the VirScan phage library [33, 34], which also encompassed peptides from a number of proteins of human epidemic and non-human CoV isolates, including MERS-CoV, SARS-CoV, as well as bat, bovine, porcine and feline isolates belonging to the alpha- and betacoronavirus genera, albeit with varying coverage of the viral peptidomes owing to the limitation in available sequence data for the latter isolates in UniProt (Supplementary Table S1). SARS-CoV-2 peptides were not included in the VirScan phage library used in our study.

We were able to obtain antibody repertoires for a total of 1399 individuals from the human cohorts described above (Supplementary Table S2). Using stringent filter criteria (Methods), we identified a total of 417 out of 2498 peptides and potential antigens from endemic HCoVs with our screen that were significantly enriched in at least three of all 1399 analyzed individuals. A total of 103 peptides from endemic HCoV were enriched in ≥ 1% of the samples and therefore considered to contain potentially immunodominant regions (Supplementary Table S3). Only 33 of the 417 peptides enriched in at least three samples shared linear sequence homology with epitopes that have previously been reported [35] (Supplementary Figure S2). To estimate number of newly identified linear B cell epitopes, we assigned each CoV-derived peptide to clusters of peptides that share ≥ 7 amino acids linear sequence identity—the estimated size of a linear B cell epitope (Methods). The enriched peptides could be assigned to 149 clusters for which at least 2 peptides share linear sequence identity of ≥ 7 amino acids (Supplementary Tables S3 and S4). Only 13 clusters also shared ≥ 7 amino acids linear sequence identity with known linear B cell epitopes. Consequently, we have identified a minimum of 136 new linear epitopes, including 25 new immunodominant linear B cell epitopes [i.e. B cell epitopes targeted in at least ≥ 1% of all individuals and not already reported in IEBD (www.iebd.org) [35] (Supplementary Tables S3)].

Next we assessed the seroprevalence of HCoV-229E, -NL63, -HKU1 and -OC43 in the three cohorts separately. To do so, we imputed species score values as described earlier [31, 33, 36] by counting the significantly enriched peptides for a given HCoV species that share less than 7 amino acids linear sequence identity. We considered an individual seropositive for any of the endemic HCoVs if the number of non-homologous peptides enriched in a given sample met our previously established species-specific cut-off value (Methods). Seroprevalence for endemic HCoVs ranged from ~4% to ~27%, depending on the species and cohort (Figure 1A). Interestingly, we found a marginal but significant negative association between age and seroprevalence of HCoV-OC43 (β = −0.175) and -NL63 (β = −0.315) (Figure 1B and 1D), as well as a marginal positive association between male gender and seroprevalence for any of the endemic HCoVs (β ≤ 0.2) (Figure 1C and 1D). The species score values (i.e. the antibody repertoire breadth for each HCoV species) did not differ substantially between seropositive individuals of our 3 cohorts (Supplementary Figure S3). However, principal component analysis revealed considerable qualitative differences in the antibody repertoires between our cohorts and in particular between pediatric and adult subjects (Figure 2A). For comparison, we also performed the same analysis on enriched peptides from other common respiratory viruses, including HRSV, HRV A, HRV B and influenza B virus. As expected, seroprevalence was considerably higher (68% to 99%) for HRSV, HRV A and HRV B, and somewhat higher (29% to 47%) for influenza B virus (Supplementary Figure S4A). However, contrary to antiviral antibody responses to endemic HCoVs, we did not find considerable variance in the antibody repertoires to other respiratory viruses when comparing age groups and cohorts (Supplementary Figure S4B). We also analyzed the enriched antigenic peptides for each endemic HCoV species separately and found that most variance in the antibody repertoires between cohorts and age groups were attributable to past infections with HCoV-HKU1 and −229E (Supplementary Figure S4C). To determine the antibody specificities responsible for most of the variance in the antiviral response to endemic HCoVs between adults and children (i.e. to identify those peptides that were significantly more or less frequently enriched when comparing adult and pediatric donors), we applied Fisher's exact test and computed odds ratios for each of the significantly enriched peptides. We found that antibody specificities in samples of pediatric study subjects predominantly targeted different antigenic regions in the S protein (mean log odds ratio = 3.35 ± 2.12), the N protein (mean log odds ratio = 2.21 ± 1.41) and diverse antigenic sites in pp1ab, whereas peptides encoding a single linear B cell epitope of pp1ab (cluster 22) appeared to be the predominant target of IgG antibodies among adult donors (mean log odds ratio = −4.7 ± 1.16) (Figure 2B and Table 1).

**Figure 1.**
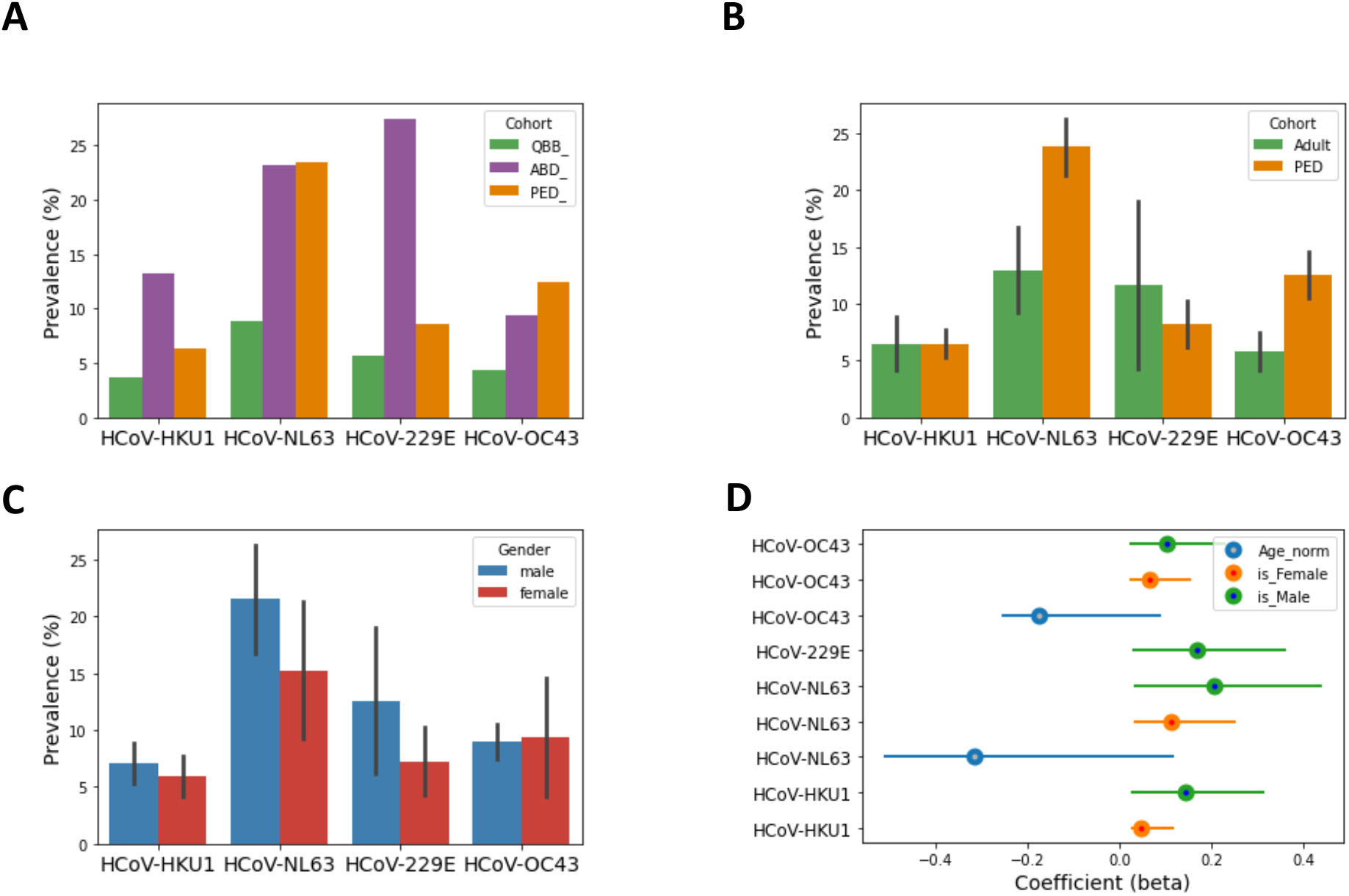
Seroprevalence of endemic HCoVs. **A-C,** Bar plots depict the seroprevalence of the four endemic HCoVs stratified by cohort (A), age group (B) and gender (C). Error bars in (B) depict gender-specific variation. Adults include adult blood bank donors and individuals of the Qatar Biobank cohort. QBB, Qatar Biobank cohort; ABD, adult (male) blood bank donors; PED, pediatric study subjects. **D**, Coefficient of association (beta) with 95% confidence interval (95% CI) of seroprevalence for each HCoV with either male gender (green), female gender (orange), or age (blue). Only features that had a P-value of association ≥ 0.001 are shown.

**Figure 2.**
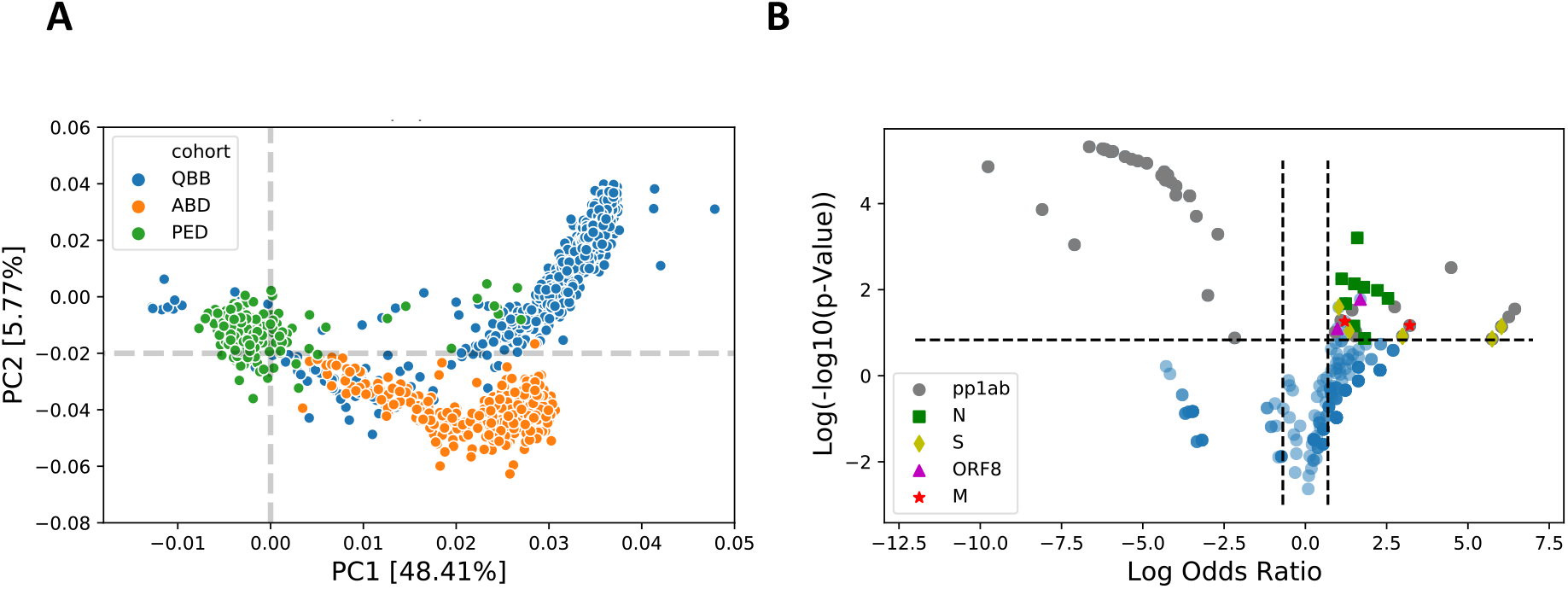
Qualitative differences in antibody repertoires between cohorts and age groups. **A,** Principal component analysis of 417 peptides from endemic HCoVs that were found to be enriched in at least 3 samples. QBB, Qatar Biobank cohort; ABD, adult (male) blood bank donors; PED, pediatric subjects. **B,** Differential enrichment analysis to determine the peptides that are either more or less frequently enriched in children versus adults (including subjects of both adult cohorts, namely QBB and ABD). We considered a peptide significantly more or less frequently enriched among children if the odds ratio (OR) was ≥ 2 or ≤ −2, respectively; and the P-value was ≤ 0.005 (Fisher’s Exact Test). pp1ab, Orf1ab replicase polyprotein; S, spike glycoprotein; M, matrix glycoprotein; N, nucleocapsid protein; ORF8, open reading frame 8 protein.

**Table 1.** List of peptides that were differentially enriched in children versus adults. Immunodominant peptides are marked in bold font.

| Entry | Start | End | Protein | Species | Log2(OR) | -Log10(P value) | Cluster |
| --- | --- | --- | --- | --- | --- | --- | --- |
| P0C6X6 | 1401 | 1456 | pp1ab | HCoV-OC43 | 6.443 | 4.717 | 136 |
| Q0ZJJ1 | 813 | 868 | pp1ab | HCoV-HKU1 | 6.257 | 3.927 | 102 |
| P0C6X2 | 7029 | 7084 | pp1ab | HCoV-HKU1 | 6.032 | 3.138 | 117 |
| Q0ZJJ1 | 225 | 280 | pp1ab | HCoV-HKU1 | 6.032 | 3.138 | 38 |
| Q5MQD0 | 365 | 420 | S | HCoV-HKU1 | 6.032 | 3.138 | 189 |
| P0C6X5 | 3753 | 3808 | pp1ab | HCoV-NL63 | 5.744 | 2.351 | 10 |
| Q0QJI4 | 337 | 392 | S | HCoV-OC43 | 5.744 | 2.351 | 98 |
| Q0ZJJ1 | 5825 | 5880 | pp1ab | HCoV-HKU1 | 5.744 | 2.351 | 9 |
| **P0C6X3** | **5489** | **5544** | **pp1ab** | **HCoV-HKU1** | **4.483** | **12.331** | **8** |
| P0C6X6 | 2381 | 2436 | pp1ab | HCoV-OC43 | 3.212 | 3.212 | 253 |
| Q6Q1R9 | 197 | 226 | M | HCoV-NL63 | 3.212 | 3.212 | 185 |
| P0C6X1 | 1373 | 1428 | pp1ab | HCoV-229E | 2.987 | 2.500 | 235 |
| Q6Q1S2 | 421 | 476 | S | HCoV-NL63 | 2.987 | 2.500 | 131 |
| Q6Q1S2 | 449 | 504 | S | HCoV-NL63 | 2.987 | 2.500 | 131 |
| P0C6X2 | 3305 | 3360 | pp1ab | HCoV-HKU1 | 2.745 | 4.957 | 13 |
| **Q6Q1R8** | **29** | **84** | **N** | **HCoV-NL63** | **2.539** | **6.042** | **49** |
| **E2DNV6** | **1** | **56** | **N** | **HCoV-NL63** | **2.217** | **7.278** | **37** |
| Q5SBN5 | 1 | 56 | N | HCoV-NL63 | 1.823 | 2.374 | 49 |
| Q6Q1R8 | 169 | 224 | N | HCoV-NL63 | 1.823 | 2.374 | 37 |
| **Q6Q1R8** | **197** | **252** | **N** | **HCoV-NL63** | **1.798** | **7.825** | **37** |
| **Q5MQC5** | **1** | **56** | **ORF8** | **HCoV-HKU1** | **1.690** | **5.895** | **41** |
| **Q6Q1R8** | **337** | **377** | **N** | **HCoV-NL63** | **1.596** | **24.578** | **132** |
| **P0C6X2** | **5461** | **5516** | **pp1ab** | **HCoV-HKU1** | **1.547** | **2.802** | **34** |
| **P0C6X5** | **4285** | **4340** | **pp1ab** | **HCoV-NL63** | **1.531** | **2.513** | **78** |
| **E2DNV6** | **29** | **84** | **N** | **HCoV-NL63** | **1.508** | **8.475** | **37** |
| **Q5SBN5** | **29** | **83** | **N** | **HCoV-NL63** | **1.492** | **3.178** | **49** |
| **Q0ZJJ1** | **1121** | **1176** | **pp1ab** | **HCoV-HKU1** | **1.437** | **4.602** | **12** |
| **P15423** | **645** | **700** | **S** | **HCoV-229E** | **1.347** | **2.822** | **16** |
| **G9G2X4** | **1** | **56** | **N** | **HCoV-HKU1** | **1.237** | **5.340** | **41** |
| **Q01455** | **197** | **230** | **M** | **HCoV-OC43** | **1.212** | **3.536** | **147** |
| **Q6Q1R8** | **309** | **364** | **N** | **HCoV-NL63** | **1.116** | **9.498** | **132** |
| **P0C6X2** | **4509** | **4564** | **pp1ab** | **HCoV-HKU1** | **1.091** | **3.625** | **2** |
| **Q0QJI4** | **757** | **812** | **S** | **HCoV-OC43** | **1.027** | **4.914** | **46** |
| **Q5MQC5** | **29** | **84** | **ORF8** | **HCoV-HKU1** | **0.984** | **2.982** | **41** |
| **P0C6X2** | **1065** | **1120** | **pp1ab** | **HCoV-HKU1** | **0.916** | **2.752** | **12** |
| **P0C6X1** | **5377** | **5432** | **pp1ab** | **HCoV-229E** | **-2.173** | **2.406** | **9** |
| **Q0ZJJ1** | **965** | **1020** | **pp1ab** | **HCoV-HKU1** | **-2.692** | **26.841** | **12** |
| **P0C6X4** | **897** | **952** | **pp1ab** | **HCoV-HKU1** | **-3.000** | **6.469** | **12** |
| **Q0ZJG7** | **925** | **980** | **pp1ab** | **HCoV-HKU1** | **-3.355** | **40.738** | **12** |
| **P0C6X4** | **925** | **980** | **pp1ab** | **HCoV-HKU1** | **-3.554** | **65.238** | **12** |
| **P0C6X3** | **925** | **980** | **pp1ab** | **HCoV-HKU1** | **-3.562** | **65.438** | **12** |
| **P0C6X4** | **953** | **1008** | **pp1ab** | **HCoV-HKU1** | **-3.977** | **81.879** | **12** |
| **P0C6X2** | **1037** | **1092** | **pp1ab** | **HCoV-HKU1** | **-3.983** | **66.553** | **12** |
| **P0C6X4** | **1009** | **1064** | **pp1ab** | **HCoV-HKU1** | **-4.132** | **89.820** | **12** |
| **Q0ZJJ1** | **1037** | **1092** | **pp1ab** | **HCoV-HKU1** | **-4.232** | **106.086** | **12** |
| **P0C6X3** | **1009** | **1064** | **pp1ab** | **HCoV-HKU1** | **-4.305** | **94.078** | **12** |
| **P0C6X2** | **951** | **1006** | **pp1ab** | **HCoV-HKU1** | **-4.339** | **114.588** | **12** |
| **P0C6X3** | **1037** | **1092** | **pp1ab** | **HCoV-HKU1** | **-4.421** | **105.026** | **12** |
| **P0C6X4** | **981** | **1036** | **pp1ab** | **HCoV-HKU1** | **-4.878** | **139.489** | **12** |
| **P0C6X2** | **925** | **980** | **pp1ab** | **HCoV-HKU1** | **-5.151** | **147.090** | **12** |
| **Q0ZJJ1** | **925** | **980** | **pp1ab** | **HCoV-HKU1** | **-5.348** | **153.250** | **12** |
| **P0C6X3** | **953** | **1008** | **pp1ab** | **HCoV-HKU1** | **-5.546** | **162.652** | **12** |
| **P0C6X3** | **951** | **1006** | **pp1ab** | **HCoV-HKU1** | **-5.924** | **183.493** | **12** |
| **Q0ZJJ1** | **1093** | **1148** | **pp1ab** | **HCoV-HKU1** | **-5.999** | **182.148** | **12** |
| **P0C6X2** | **949** | **1004** | **pp1ab** | **HCoV-HKU1** | **-6.155** | **192.846** | **12** |
| **Q0ZJJ1** | **953** | **1008** | **pp1ab** | **HCoV-HKU1** | **-6.176** | **190.949** | **12** |
| **P0C6X2** | **953** | **1008** | **pp1ab** | **HCoV-HKU1** | **-6.238** | **195.471** | **12** |
| **Q0ZJJ1** | **979** | **1034** | **pp1ab** | **HCoV-HKU1** | **-6.647** | **204.612** | **12** |
| **P0C6X4** | **4761** | **4816** | **pp1ab** | **HCoV-HKU1** | **-7.102** | **20.977** | **36** |
| **Q0ZJG7** | **4985** | **5040** | **pp1ab** | **HCoV-HKU1** | **-8.096** | **47.573** | **21** |
| **P0C6X1** | **337** | **392** | **pp1ab** | **HCoV-229E** | **-9.759** | **128.638** | **219** |

Intriguingly, multiple sequence alignments of frequently enriched peptides with the full-length proteins of various CoVs revealed that antibody specificities predominantly found in pediatric study subjects target immunodominant epitopes that encode functionally important and highly conserved regions of the structural proteins. These included regions in the S_1_ subunit of the S protein which are important for receptor binding (Figure 3A-C and 3F) [37–40], as well as the regions resembling the proteolytic cleavage sites and fusion peptide of the S_2_ subunit (Figures 3A, 3B, 3D-F). Of note, the immunodominant region spanning the furin-like S2’ cleavage site in the S_2_ subunit resembles one of the most conserved regions of the S protein, both in amino acid sequence (R↓SA[I/L]ED[I/L]LF) (Figure 3E) and in protein structure, as it forms an accessible alpha-helix within the fusion peptide region (Figure 3F and Supplementary Figure S6) [41]. Moreover, we identified potential antibody binding sites in the N-terminal RNA binding domain, serine-rich region and in the C-terminal dimerization domain of the N protein (Figures 4A and 4B). Although the predicted antibody binding sites in the N-terminal RNA-binding domain and the C-terminal dimerization domain of the N protein appeared to be less conserved between different species in the primary amino acid sequence (Figure 4C and 4D), both domains are structurally conserved in the regions that we found to be immunodominant (Figure 4E and 4F). We also found that antibodies in children targeted more frequently the C-terminal domain of the M protein (Supplementary Figure S5C and Table 1) and the small accessory Orf8 protein (also known as N2) of HCoV-HKU1 (Table 1). Although Orf8 and N share the same coding sequence in the viral RNA genome, the reading frame is different and the amino acid sequences not homologous. On the contrary, antibody specificities predominantly found in adults primarily targeted a region of the pp1ab that is specific to HCoV-HKU1 and contains an acidic tandem repeat (NDDE[D/H]VVTGD) which is located upstream of the papain-like protease 1 domain (Supplementary Figure S5D).

**Figure 3.**
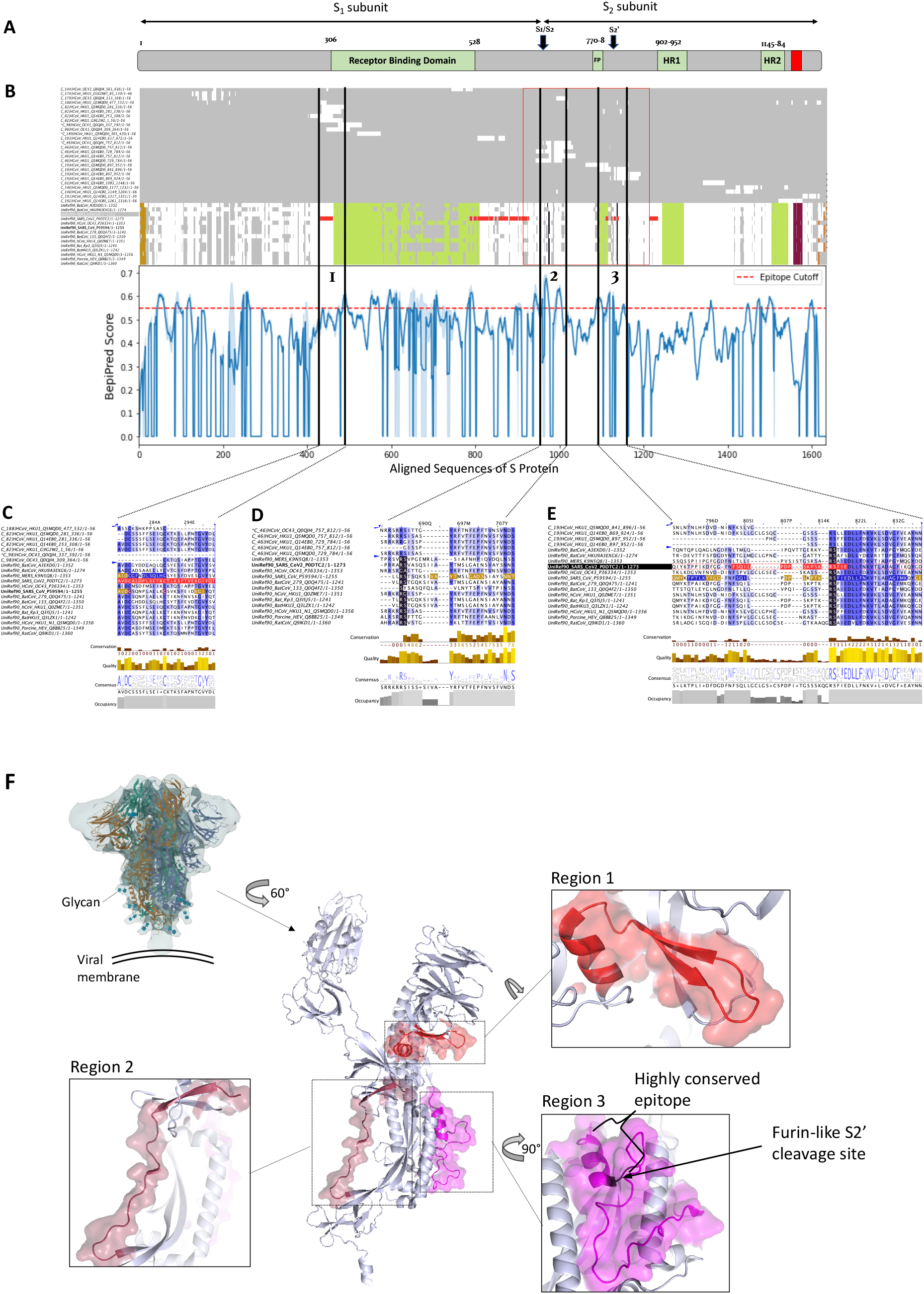
Antigenic regions and predicted antibody binding sites of the spike (S) protein. **A,** Schematic representation of the S protein of SARS-CoV (strain Tor2). Proteolytic cleavage sites of the S protein are marked with arrows. **B,** Overview of a multiple sequence alignment of immunodominant peptides with the full-length protein sequences of various alpha- and beta-CoVs (top). The mean score (blue line) and standard deviations (shaded) for the residue-wise prediction of linear B cell epitopes of aligned endemic HCoVs (bottom). Row labels (left) indicate the cluster and sequence identifier, start and end position of the peptide and length of the aligned sequence. Peptides for which differential enrichment between children and adults was statistically significant (P-value ≤ 0.05, Fisher’s exact test) and odds ratios were ≥ 2 are indicated with a “*”. The protein domains and boundaries shown in the schematic (A) have been marked in different colors using UniProt annotation and JalView features. Amino acid sequences of previously predicted immunodominant linear SARS-CoV-2 B cell epitopes [43] are highlighted in pink. For the BepiPred score (bottom), a score cutoff of 0.55 has been marked with a dashed red line to indicate regions that are predicted to be potential B cell epitopes. **C-E,** Selected regions of the multiple sequence alignment encompassing regions 1, 2 and 3 as shown in (B). Proteolytic cleavage sites of the S protein are highlighted in black. The full sequence alignment is shown in Supplementary Figure S5A. **F,** Ribbon diagram of the SARS-CoV-2 S trimer with carbohydrate residues shown (top left) and monomer without carbohydrate residues (middle) in the prefusion conformation (PDB id: 6VXX, chain A) [41]; regions 1, 2 and 3 are shown enlarged. FP, fusion peptide; HR1 and HR2, heptad repeat 1 and 2.

**Figure 4.**
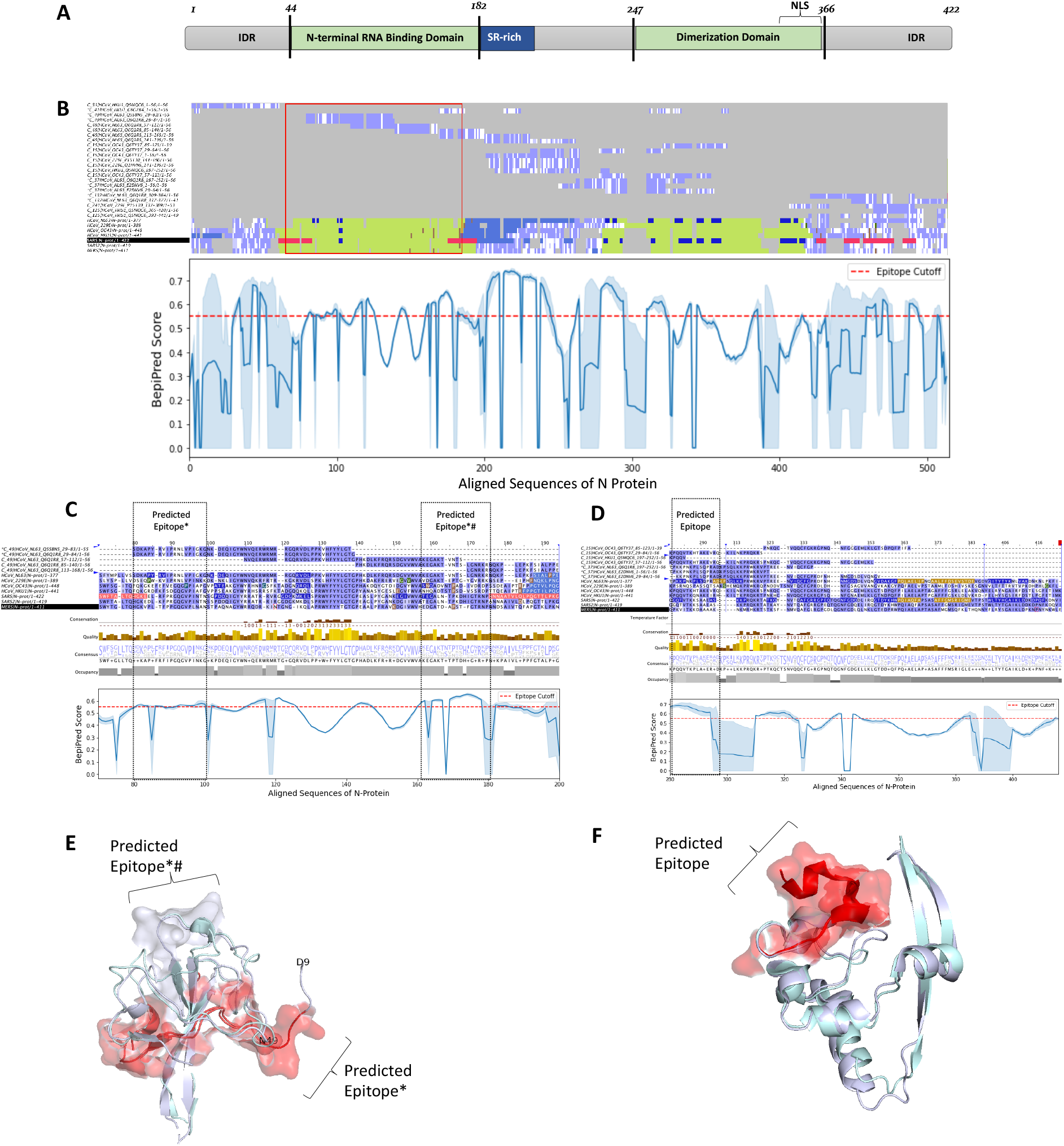
Antigenic regions and predicted antibody binding sites of the nucleocapsid (N) protein. **A,** Schematic representations of the N protein of SARS-CoV (strain Tor2). SR-rich, serine-rich; NLS, Predicted nuclear localization sequence; IDR, intrinsically disordered region. **B,** Overview of a multiple sequence alignment of immunodominant peptides with the full-length protein sequences of various alpha- and beta-CoVs (top) and mean score values for the prediction of linear B cell epitopes among endemic HCoVs (bottom). Row labels (left) indicate the cluster and sequence identifier, start and end position of the peptide and length of the aligned sequence. Peptides for which differential enrichment between children and adults was statistically significant (P-value ≤ 0.005, Fisher’s exact test) and odds ratios were ≥ 2 are indicated with a “*”. The protein domains of full-length reference sequences and boundaries shown in the schematic (A) have been marked in green. Amino acid sequences of previously predicted immunodominant SARS-CoV B cell epitopes [43] are highlighted in pink color. For the BepiPred score (bottom), a score cutoff of 0.55 has been marked with a dashed red line to indicate regions that are predicted to be potential B cell epitopes. **C, D,** Selected regions of the multiple sequence alignment encompassing the N-terminal RNA-binding domain (C) and C-terminal self-assembly domain (D). The full multiple sequence alignment is shown in Supplementary Figure S5B. **E,** Super-imposed ribbon structure of the N-terminal RNA-binding domain from HCoV-NL63 (PDB: 5N4k, chain A) and that from SARS-CoV-2 (PDB: 6M3M, chain A) (root-mean-square deviation [rmsd] = 0.7 Ångström). **F,** Super-imposed ribbon structure of the C-terminal self assembly domain of proteins from HCoV- NL63 (pdbId: 5EPW, chain A) and that of SARS-CoV-2 (pdbID: 6WJI, chain A) (rmsd = 0.91 Ångström). *, Predicted epitopes in peptides for which differential enrichment between children and adults was statistically significant (P-value ≤ 0.005, Fisher’s exact test) and odds ratios were ≥ 2; #, Predicted immunodominant epitopes.

Given the high degree of sequence conservation among some of the immunodominant regions in proteins of endemic HCoVs we have identified, we also explored the extent to which antibody specificities to these regions cross-react with peptides from epidemic CoVs and non-human CoV isolates. For this purpose, we assessed the enrichment of peptides derived from SARS-CoV, MERS-CoV as well as bovine, porcine, bat and feline CoV isolates (Supplementary Table S1) applying the same approach and stringent filter criteria as described above, for peptides of endemic HCoVs. Indeed, we identified several S protein- and N protein-derived peptides from epidemic CoVs or non-human isolates that were significantly enriched in our PhIP-Seq assay, which share sequence similarity with peptides from HCoVs (Figure 5). As expected based on the results from multiple sequence alignments described above, antibody specificities targeting the highly conserved amino acid motif (RSA[I/L]ED[I/L]LF) spanning the furin-like S2’ cleavage site of the S protein were broadly cross-reactive to several orthologous peptides from MERS-CoV, SARS-CoV and non-human CoV isolates (Figure 5A, Region 3). Cross-reactivity of antibodies targeting other functionally important but less conserved regions of the S protein also appeared to be more restricted (Figure 5A, Region 1 and 2). Antibody specificities targeting the N protein also showed considerable cross-reactivity with peptides from MERS-CoV, SARS-CoV and non-human CoV isolates. However, the latter cross-reactive antibodies mainly targeted regions rich in serine and arginine residues, with low-complexity sequences and very limited structural conservation, particularly an intrinsically disordered region (IDR) at the N terminus of the N protein (Figure 5B and Supplementary Figure 7) that lacks a tertiary structure [42]. We also detected cross-reactive antibodies targeting the serine- and arginine-rich linker region of the N protein; however, cross-reactivity to this region was largely restricted to peptides derived from non-human CoV isolates of domestic animals (Figure 5B) which are more closely related to HCoV-OC43 [6].

**Figure 5.**
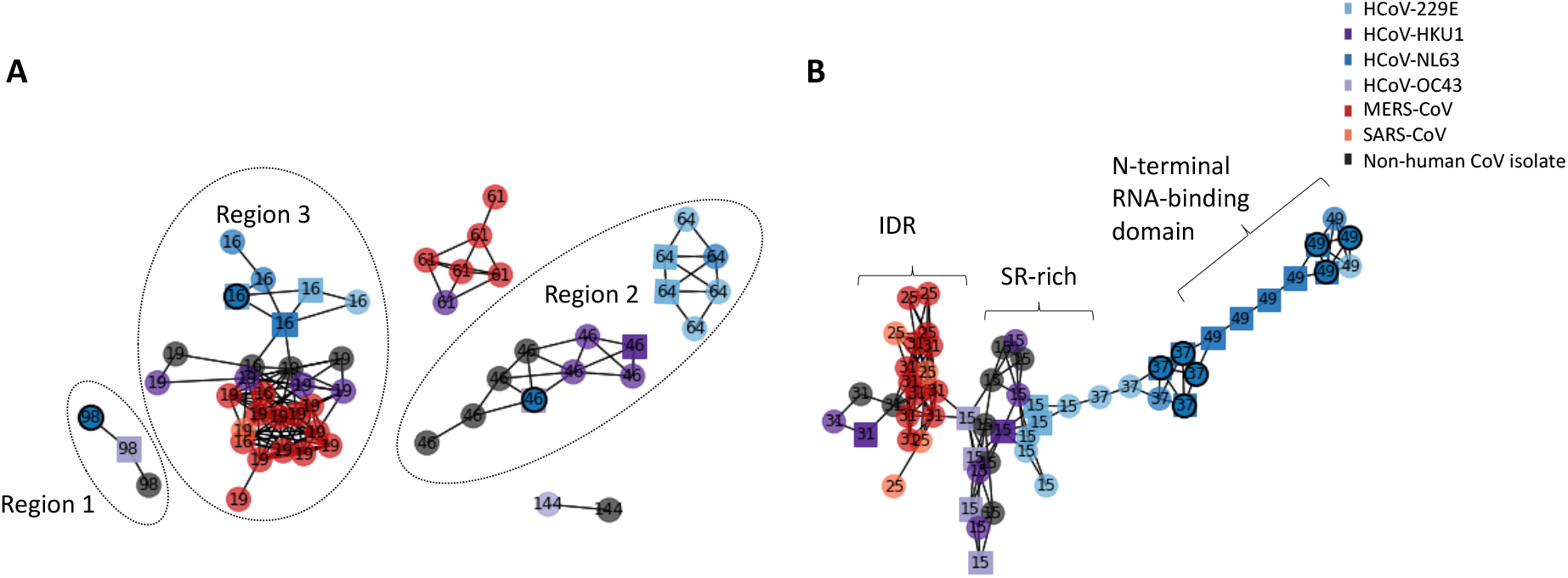
Network representation of enriched peptides from structural proteins targeted by cross-reactive antibodies. **A,** Network representation of enriched spike (S) protein-derived peptides. **B,** Network representation of enriched nucleoprotein (N)-derived peptides. Each node represents an enriched peptide and the color indicates the species. Edges indicate ≥ seven amino acids linear sequence identity between two nodes (i.e. peptides), the estimated size of a linear B cell epitope. Only networks of peptides derived from at least two different species are shown. Labels indicate the cluster number to which each peptide has been assigned. Nodes are represented as spheres if the peptide had been frequently enriched. Nodes marked with a black circle indicate peptides for which differential enrichment between children and adults was statistically significant (P-value ≤ 0.005, Fisher’s exact test) and odds ratios were ≥ 2. SR-rich, serine- and arginine-rich motive; IDR, intrinsically disordered region.

Finally, we assessed plasma samples of a previously healthy 49-year-old female adult patient who suffered from severe ARDS from SARS-CoV-2 infection, requiring prolonged hospitalization and intensive care. A first sample was obtained 25 days after onset of symptoms and 18 days after ICU admission. A second sample was obtained one month later, one week after discharge. We compared the antibody profiles at both time points to samples obtained from uninfected family members of the patient, as well as an age- and gender-matched unrelated control. In agreement with what we found in some subjects with a history of endemic HCoV infection, the patient with severe COVID-19 had detectable antibodies that cross-reacted with peptides from SARS-CoV and non-human CoVs encoding the furin-like S2’ cleavage site and heptad repeat 2 region of the S protein, peptides from the C-terminal region of the HCoV-HKU1 N protein downstream of the dimerization domain, as well as two antigenic sites of the MERS-CoV pp1a (Supplementary Figure S8).

## Discussion

Our comprehensive and systematic screen for antiviral antibody repertoires across individuals in three human cohorts revealed a large number of peptides with novel linear B cell epitopes in several proteins of endemic HCoVs. This is not surprising given that epidemic CoVs, and in particular SARS-CoV, have been the primary focus of previous immunological and epitope screening studies [35, 43]. Information about the targets of immune responses to CoVs across different species provides a valuable resource for the prediction of candidate targets of newly emerging CoVs, as recently shown by Grifoni *et al.* [43]. The authors were able to identify *a priori,* several specific regions of the S, M and N proteins of SARS-CoV-2 on the basis of sequence homology to the SARS-CoV virus which are orthologous to several of the immunodominant regions of endemic HCoVs we identified. We detected antibodies against the structural S, N, M and Orf8 proteins, as well as the non-structural pp1ab polyprotein of HCoVs; the latter resembling the precursor for the large viral replicase complex [44]. Interestingly, in another independent study, Grifoni *et al.* [28] recently reported similarly broad T cell responses in COVID-19 patients by employing an analogous screen for T cell epitopes of SARS-CoV-2 proteins using peptide ‘‘megapools’’ in combination with *ex vivo* T cell assays.

Surprisingly, circulating IgG antibodies in children appear to be differentially targeting structural and non-structural proteins of HCoVs in comparison to adults (Figure 2). Whereas antibody specificities more frequently found in samples of pediatric subjects targeted structural proteins such as the S, N and M proteins, in adult donors, a region of the non-structural polyprotein pp1ab containing an acidic tandem repeat (NDDE[D/H]VVTGD) in HCoV-HKU1 appeared to be the predominant target of IgG antibodies. The latter polyprotein is post-translationally processed into up to 16 subunits that form a large viral replicase complex; however, the function of the acidic tandem repeat and its role in pathogenesis remains unknown [44]. This qualitative difference in the antibody repertoires of adult versus pediatric subjects appeared to be a specific characteristic of natural HCoV infection, as we did not find the same degree of variance in the antibody repertoires specific to other common respiratory viruses when comparing our cohorts and different age groups (Supplementary Figure S3). We speculate that the qualitative differences in antibody repertoires of adult versus children reflect a higher frequency and/or more recent exposures of children to seasonal coronaviruses than adults, coupled with the rapid decay of circulating anti-CoV antibodies that target the structural proteins of the virions. Further studies will be needed to fully understand the dynamics of antibody responses to endemic HCoVs.

Evidence for the transient and dynamic nature of humoral immunity to endemic HCoVs has been provided by number of human serological studies, although many of them were conducted with only a limited number of subjects, or only for selected species, and a variety of antibody detection methods were used that are not readily comparable [45–52]. Nevertheless, evidence suggests that a sizable proportion of children experience primary infection with endemic HCoVs during their first year of life and nearly all children have encountered at least one of the endemic HCoVs before two years of age, indicating that first exposure to endemic HCoVs occurs very early in life, similar to other common respiratory viruses such as HRSV or HRVs [46, 50, 51]. However, reported seroprevalence rates in older children and adults vary greatly depending on a variety of factors, including age and viral species. There is a general trend indicating that humoral immunity from primary infection with endemic HCoVs wanes quickly and that antibodies detected in older children and adults are rather a consequence of more recent re-infections [45–49]. We estimated the seroprevalence of antibodies to endemic HCoVs to range between ~4% and ~27% depending on the species and cohort (Figure 1A-C). Given that endemic HCoV infections are common and usually acquired during early childhood [46, 47, 50, 51], it is likely that not only the adult subjects, but also many (if not all) of the children aged 7 to 15 years that were assessed in our study, have already experienced multiple infections with endemic HCoVs in their lifetime. Therefore, our estimated seroprevalence rates likely reflect the complex dynamics between rates of (re-)infection and waning humoral immunity over time. In agreement with this notion, age was negatively associated with seroprevalence in our study, suggesting that the duration of immunity in response to natural infection with endemic HCoVs and/or rates of re-infection reduce with increasing age. The dynamics of humoral immunity from past CoV infections is best described in studies of MERS and SARS patients. Although limited in sample size, these studies have shown that antibody titers in all previously infected individuals decline relatively quickly to minimal detectable levels over a period of 2 to 3 years and that patients who suffered from more severe disease had higher and longer-lasting total binding antibody titers and neutralizing titers [52]. There is also evidence that symptomatic COVID-19 patients mount robust antibody responses that wane very quickly over a period of 6 months [53]. However, the same study suggeted that SARS-CoV-2-specific memory B cells may be sustained over a longer period of time [53]. Indeed, most acute virus infections induce some level of protective and long-term immunity, albeit through a variety of mechanisms that are not necessarily the same for each pathogen and may even differ between hosts due to a variety of factors, including simultaneous viral coinfection [54, 55]. We also found a marginal but significant positive association between seroprevalence of endemic HCoVs and male gender, which is consistent with an earlier report by Gaunt *et al.* [5].

Despite the variable degree of sequence conservation between different CoV species, the results of our systematic antibody screen highlight that the structural proteins of the virions share common antigenic sites. Indeed, several of the immunodominant regions we have identified experimentally in the structural proteins of endemic HCoVs are orthologous to the regions thought to be immunodominant targets for immune responses to SARS-CoV-2 [43] (Supplementary Figure S8), including two linear epitopes on the SARS-CoV-2 S protein that elicit potent neutralizing antibodies in COVID-19 patients [56]. Importantly, antigenic regions that we found to be immunodominant in our study (i.e. enriched in ≥ 1% of all samples), as well as those corresponding to peptides for which enrichment was strongly and significantly associated with pediatric subjects (odds ratio ≥ 2; P-value ≤ 0.005, Fisher’s exact test), mapped to functionally important regions of the structural CoV proteins. These included regions for receptor binding and the proteolytic cleavage sites of the S protein, as well as the N-terminal RNA-binding and C-terminal dimerization domains of the N protein, which have been shown to be critical for virus attachment and entry, cell-to-cell fusion and virus replication, respectively [42, 57–61]. The region of the S_1_ subunit responsible for receptor binding differs considerably between CoV species, which utilize different domains and host cell receptors, and consequently differ in their tissue tropism [38–41, 47, 62]. However, the S_2_ subunit resembling the fusion machinery is more conserved, both structurally and in amino acid sequence [39, 63] (Supplementary Figure S6). Indeed, we identified an immunodominant and highly conserved linear epitope immediately downstream of the furin-like S2’ cleavage site of the S protein (R↓SA[I/L]ED[I/L]LF) that likely resembles the fusion peptide, although its precise location has been disputed [64]. The same antigenic site has recently been found on the SARS-CoV-2 S protein to elicit neutralizing antibodies in COVID-19 patients [56]. The high degree of amino acid sequence and conformational conservation of the alpha-helical region immediately adjacent to the S2’ cleavage site (Figures 3E and 3F) likely explains why antibodies targeting this region also cross-reacted with orthologous peptides of related CoVs in our study, including that of MERS-CoV, SARS-CoV and non-human isolates, further supporting our overall hypothesis and the important role of this particular region as a pan-CoV target site [41, 56]. It is therefore tempting to speculate that at least in some individuals, past infections with endemic HCoVs have elicited cross-reactive antibodies and/or led to the generation of longer-lived memory B cells with specific reactivity to this linear epitope, which may provide cross-protection against MERS or COVID-19. This may be the case particularly among children which are generally less likely to experience severe disease outcomes from infection with epidemic CoVs [17–19, 21]. This will require further investigation.

Broadly cross-reactive antibody responses are also known for other enveloped RNA viruses, which may positively or negatively affect subsequent infection or vaccination. Flaviviruses for example, are antigenically related and broadly flavivirus cross-reactive antibodies from previous yellow fever vaccination has been shown to impair and modulate immune responses to tick-borne encephalitis vaccination [65]. Similarly, immune history has been shown to profoundly affect protective B cell responses to influenza [66]. Since we detected pan-CoV cross-reactive antibodies less frequently in plasma samples from adult donors, our results argue against a strong therapeutic benefit of intravenous immunoglobulin products to control the spread of COVID-19 disease [67]. In this context, it should be noted that large-scale antibody screening by PhIP-Seq may frequently fail to detect conformational and post-translationally modified B cell epitopes [29]. Nontheless, we found anti-CoV antibodies in plasma of a COVID-19 patient after prolonged hospitalization and intensive care that targeted largely the same structurally conserved and functionally important regions of the viral N and S proteins (Supplementary Figure S8) as those that we detected in a sizable proportion of children, including antibodies binding to the highly conserved motif and furin-like S2’ cleavage site (R↓SA[I/L]ED[I/L]LF), which provides further evidence for the clinical benefit of using convalescent plasma for the prevention and treatment of COVID-19 [68–70]. Our findings may also have important implications for the development of prophylactic or therapeutic monoclonal antibodies and vaccine design in the context of COVID-19 [43, 71].

## Methods

### Study design and samples

We performed a retrospective analysis of deidentified or coded plasma and serum samples collected from three different human cohorts, namely: i) 400 healthy male adult blood donors (ABD) of a blood bank in Qatar with diverse ethnic background and nationality (Supplementary Figure S1A); ii) 800 adult male and female Qatari nationals and long-term residents of Qatar who are participating in a national cohort study— the Qatar Biobank (QBB)—and who represent the general local population in the State of Qatar [32]; and iii) 231 pediatric subjects with Qatari nationality who were admitted to, or visited outpatient clinics of, Sidra Medicine. Leftover plasma samples from healthy blood bank donors were collected from 2012 to 2016, de-identified and stored at −80°C. For the purpose of this study, specimen from male Qatari nationals 19 to 66 years of age (Supplementary Table S1) were selected from a larger blood donor cohort including 5983 individuals, and then age-matched male donors with other nationalities were randomly selected (Supplementary Figure S1). Samples from female blood bank donors were excluded because they were largely underrepresented in the blood bank donor cohort. We also excluded samples for which age, gender or nationality information was lacking. Serum samples from the QBB cohort were collected from 2012 to 2017 and were randomly selected samples from the first 3000 individuals taking part in a longitudinal cohort study as described previously [32]. Plasma samples from pediatric patients were selected from leftovers of samples processed in the clinical chemistry labs of Sidra Medicine, a tertiary care hospital for children and women in Doha, Qatar, over a period of several months from September to November 2019. In order to select appropriate pediatric samples, electronic medical records were queried using Discern analytics to identify blood samples from Qatari nationals aged 7 to 15 years for whom basic metabolic panel (BMP) and comprehensive metabolic panel (CMP) testing was done in the previous week. Samples from oncology patients, patients requiring complex care and those in intensive care units, as well as samples from patients with chronic diseases, samples with no centile data and samples from patients who were underweight (centile <5%) or overweight were excluded. However, we included obese patients in our analysis, since a considerable proportion of Qatari nationals are overweight. The COVID-19 patient assessed in this study was a previously healthy female Belgian national with autosomal recessive IRF7 deficiency who developed ARDS following SARS-CoV-2 infection at the age of 49 [72]. For comparison, we also assessed unexposed family members, including the father, mother, brother (heterozygous carriers) and wild-type sister, as well as an unrelated age- and gender-matched healthy control. The human subject research described here had been approved by the institutional research ethics boards of Sidra Medicine, Qatar Biobank, INSERM, and Erasme Hospital.

### Phage Immunoprecipitation-Sequencing (PhIP-Seq)

Large scale serological profiling of the antiviral IgG repertoires in the individual serum or plasma samples was performed as described by Xu *el al.* [31]. Each serum or plasma sample was tested in duplicate and samples were analyzed in batches with up to 96 samples per batch. Only samples that satisfied a minimum read count of 1*10^6^ as well as a Pearson correlation coefficient of ≥ 0.7 in the two technical repeats were considered for downstream analysis. Data from thirty individuals of the ABD cohort and two individuals of the QBB cohort were excluded from the downstream analysis due to insufficient sequencing read depth, low sequencing data quality or because one of two technical replicates had failed (data not shown).

### Peptide enrichment analysis

To filter for enriched peptides, we first imputed −log10(P) values as described previously [31, 33, 36] by fitting a zero-inflated generalized Poisson model to the distribution of output counts and regressed the parameters for each peptide sequence based on the input read count. We considered a peptide enriched if it passed a reproducibility threshold of 2.3 [−log10(P)] in two technical sample replicates. To remove sporadic hits, we then filtered for antibody specificities to CoV peptides that were found to be enriched in at least three of all 1399 subjects assayed and analyzed in this study. We computed species-specific significance cut-off values to estimate minimum number of enriched, non-homologous peptides required in order to consider a sample as seropositive using a generalized linear model and in-house serological (ELISA) data from pooled samples that were tested positive for various viruses. We then computed virus score values as described by Xu et al [31] by counting enriched, non-homologous peptides for a given species and then adjusted these score values by dividing them with the estimated score cutoff. For the purpose of this study and under consideration of the seroprevalence of endemic HCoVs in our three cohorts, we considered a peptide to be immunodominant if it was enriched in ≥ 1% of all 1399 subjects assayed and analyzed in this study.

### Association studies and differential enrichment analysis

We applied a generalized linear model to test for associations between the HCoV species-specific adjusted score values, gender and age. We considered an association to be significant if the P-value was ≤ 0.001. We examined the frequency distribution of enriched peptides among samples of the different age groups (PED versus ABD + QBB) by estimating odds ratios (OR) and associated P-values using Fisher’s exact test. Peptides that satisfied both significance (P-value ≤ 0.005) and magnitude criteria (|log(OR)| ≥ log(2)) were considered to be differentially enriched. Positive log(OR) values indicated more frequent peptide enrichment among pediatric study subjects, whereas negative log(OR) values indicated more frequent peptide enrichment among adult subjects.

### Clustering of peptides for shared linear B cell epitopes

To estimate the minimum number of linear B cell epitopes among the enriched peptides, we built a pairwise distance matrix that captured the maximum size of linear sequence identity of amino acids (d_i,j_) between all enriched peptides. Groups of peptides that shared ≥ seven amino acid linear sequence identity (d_i,j_ ≥ 7) were assigned to a cluster. Peptides of a given cluster were considered to share a linear B cell epitope (Supplementary Figure S7).

### Software

Open source Python modules with in-house scripts were used to test for associations (statsmodel v0.11), to filter differentially enrichment peptides and to perform different statistical tests (sklearn v0.23, scipy v1.14.1). Multiple sequence alignments were done using the MAFFT [73, 74] via EMBL-EBI’s web services and Java Alignment Viewer (Jalview) for visualization [75]. Residue-wise linear B cell epitopes were predicted using BepiPred-2.0 [76]. Protein structures graphics were generated using PyMOL (Schrödinger).

## Author contribution statement

TK & NM conceived the original idea, designed the models and the computational framework of the study, analyzed the data and wrote manuscript. MR, FA, SH, QZ, PB, ZL, EJ, VB, AC, HY, MS, LA and JLC planned or performed the experiments. GN, AS, IV, JCG, GS, IM, FH, IM and MH contributed samples and data. MR, JLC and MH provided critical feedback to this version of the manuscript. All authors have seen and approved the manuscript. It has not been accepted or published elsewhere.

## Acknowledgement

We would like to thank the Qatar Biobank (QBB) management and staff, in particular Dr. Nahla Afifi and Elizabeth Jose, for their time and effort allowing us to access and analyze samples and data from the Qatar Biobank, and Dr. Stephan Elledge (Brigham and Women’s Hospital and Harvard Medical School, Boston, MA) for kindly providing the VirScan phage library used in this study. We would also like to thank Patrick Tang, Mohammed Yousuf Karim, Evonne Chin-Smith and Damien Chaussabel (Sidra Medicine) for critical reading of the manuscript. This work was supported in part by a grant from the Qatar National Research Fund (PPM1-1220-150017) and funds from Sidra Medicine.

## Potential competing interests

None of the authors has any competing interests to declare.

## Notes

### Competing Interest Statement

The authors have declared no competing interest.

### Summary of Updates

Result section (lines 249-260) and discussion updated; Author list updated; Supplemental files updated.

## References

1. Hamre, D. and J.J. Procknow, A New Virus Isolated from the Human Respiratory Tract. Proceedings of the Society for Experimental Biology and Medicine, 1966. 121(1): p. 190–193.

2. McIntosh, K., et al., Recovery in tracheal organ cultures of novel viruses from patients with respiratory disease. Proceedings of the National Academy of Sciences, 1967. 57(4): p. 933–940.

3. van der Hoek, L., et al., Identification of a new human coronavirus. Nature Medicine, 2004. 10(4): p. 368–373.

4. Woo, P.C.Y., et al., Characterization and Complete Genome Sequence of a Novel Coronavirus, Coronavirus HKU1, from Patients with Pneumonia. Journal of Virology, 2005. 79(2): p. 884–895.

5. Gaunt, E.R., et al., Epidemiology and clinical presentations of the four human coronaviruses 229E, HKU1, NL63, and OC43 detected over 3 years using a novel multiplex real-time PCR method. J Clin Microbiol, 2010. 48(8): p. 2940–7.

6. Corman, V.M., et al., Hosts and Sources of Endemic Human Coronaviruses. Adv Virus Res, 2018. 100: p. 163–188.

7. Kristoffersen, A.W., et al., Coronavirus causes lower respiratory tract infections less frequently than RSV in hospitalized Norwegian children. Pediatr Infect Dis J, 2011. 30(4): p. 279–83.

8. Drosten, C., et al., Identification of a novel coronavirus in patients with severe acute respiratory syndrome. N Engl J Med, 2003. 348(20): p. 1967–76.

9. Zaki, A.M., et al., Isolation of a Novel Coronavirus from a Man with Pneumonia in Saudi Arabia. New England Journal of Medicine, 2012. 367(19): p. 1814–1820.

10. Zhu, N., et al., A Novel Coronavirus from Patients with Pneumonia in China, 2019. New England Journal of Medicine, 2020. 382(8): p. 727–733.

11. Dong, E., H. Du, and L. Gardner, An interactive web-based dashboard to track COVID-19 in real time. The Lancet Infectious Diseases, 2020.

12. de Wit, E., et al., SARS and MERS: recent insights into emerging coronaviruses. Nat Rev Microbiol, 2016. 14(8): p. 523–34.

13. Severe Outcomes Among Patients with Coronavirus Disease 2019 (COVID-19) — United States, February 12–March 16, 2020. MMWR Morb Mortal Wkly Rep, 2020. 69: p. 343–346.

14. Li, X. and X. Ma, Acute respiratory failure in COVID-19: is it “typical” ARDS? Critical Care, 2020. 24(1): p. 198.

15. Vijgen, L., et al., Evolutionary History of the Closely Related Group 2 Coronaviruses: Porcine Hemagglutinating Encephalomyelitis Virus, Bovine Coronavirus, and Human Coronavirus OC43. Journal of Virology, 2006. 80(14): p. 7270–7274.

16. Young, B.E., et al., Epidemiologic Features and Clinical Course of Patients Infected With SARS-CoV-2 in Singapore. JAMA, 2020.

17. Dong, Y., et al., Epidemiology of COVID-19 Among Children in China. Pediatrics, 2020: p. e20200702.

18. Zimmermann, P. and N. Curtis, Coronavirus Infections in Children Including COVID-19: An Overview of the Epidemiology, Clinical Features, Diagnosis, Treatment and Prevention Options in Children. The Pediatric Infectious Disease Journal, 2020. 39(5): p. 355–368.

19. Al-Tawfiq, J.A., R.F. Kattan, and Z.A. Memish, Middle East respiratory syndrome coronavirus disease is rare in children: An update from Saudi Arabia. World J Clin Pediatr, 2016. 5(4): p. 391–396.

20. Zhang, J., et al., Changes in contact patterns shape the dynamics of the COVID-19 outbreak in China. Science, 2020: p. eabb8001.

21. Davies, N.G., et al., Age-dependent effects in the transmission and control of COVID-19 epidemics. Nature Medicine, 2020.

22. Guo, L., et al., Profiling Early Humoral Response to Diagnose Novel Coronavirus Disease (COVID-19). Clinical Infectious Diseases, 2020.

23. Okba, N.M.A., et al., Severe Acute Respiratory Syndrome Coronavirus 2-Specific Antibody Responses in Coronavirus Disease 2019 Patients. Emerg Infect Dis, 2020. 26(7).

24. Yongchen, Z., et al., Different longitudinal patterns of nucleic acid and serology testing results based on disease severity of COVID-19 patients. Emerg Microbes Infect, 2020. 9(1): p. 833–836.

25. Khan, S., et al., Analysis of Serologic Cross-Reactivity Between Common Human Coronaviruses and SARS-CoV-2 Using Coronavirus Antigen Microarray. bioRxiv, 2020: p. 2020.03.24.006544.

26. Smatti, M.K., A.A. Al Thani, and H.M. Yassine, Viral-Induced Enhanced Disease Illness. Front Microbiol, 2018. 9: p. 2991.

27. Braun, J., et al., SARS-CoV-2-reactive T cells in healthy donors and patients with COVID-19. Nature, 2020.

28. Grifoni, A., et al., Targets of T Cell Responses to SARS-CoV-2 Coronavirus in Humans with COVID-19 Disease and Unexposed Individuals. Cell, 2020.

29. Mohan, D., et al., Publisher Correction: PhIP-Seq characterization of serum antibodies using oligonucleotide-encoded peptidomes. Nat Protoc, 2019. 14(8): p. 2596.

30. Mohan, D., et al., PhIP-Seq characterization of serum antibodies using oligonucleotide-encoded peptidomes. Nat Protoc, 2018. 13(9): p. 1958–1978.

31. Xu, G.J., et al., Viral immunology. Comprehensive serological profiling of human populations using a synthetic human virome. Science, 2015. 348(6239): p. aaa0698.

32. Al Kuwari, H., et al., The Qatar Biobank: background and methods. BMC Public Health, 2015. 15: p. 1208.

33. Mina, M.J., et al., Measles virus infection diminishes preexisting antibodies that offer protection from other pathogens. Science, 2019. 366(6465): p. 599–606.

34. Drutman, S.B., et al., Fatal Cytomegalovirus Infection in an Adult with Inherited NOS2 Deficiency. N Engl J Med, 2020. 382(5): p. 437–445.

35. Vita, R., et al., The Immune Epitope Database (IEDB): 2018 update. Nucleic Acids Res, 2019. 47(D1): p. D339–D343.

36. Pou, C., et al., The repertoire of maternal anti-viral antibodies in human newborns. Nat Med, 2019. 25(4): p. 591–596.

37. Tortorici, M.A., et al., Structural basis for human coronavirus attachment to sialic acid receptors. Nat Struct Mol Biol, 2019. 26(6): p. 481–489.

38. Tortorici, M.A. and D. Veesler, Structural insights into coronavirus entry. Adv Virus Res, 2019. 105: p. 93–116.

39. Rey, F.A. and S.M. Lok, Common Features of Enveloped Viruses and Implications for Immunogen Design for Next-Generation Vaccines. Cell, 2018. 172(6): p. 1319–1334.

40. Hulswit, R.J.G., et al., Human coronaviruses OC43 and HKU1 bind to 9-<em>O</em>-acetylated sialic acids via a conserved receptor-binding site in spike protein domain A. Proceedings of the National Academy of Sciences, 2019. 116(7): p. 2681–2690.

41. Walls, A.C., et al., Structure, Function, and Antigenicity of the SARS-CoV-2 Spike Glycoprotein. Cell, 2020. 181(2): p. 281–292 e6.

42. McBride, R., M. van Zyl, and B.C. Fielding, The coronavirus nucleocapsid is a multifunctional protein. Viruses, 2014. 6(8): p. 2991–3018.

43. Grifoni, A., et al., A Sequence Homology and Bioinformatic Approach Can Predict Candidate Targets for Immune Responses to SARS-CoV-2. Cell Host Microbe, 2020. 27(4): p. 671–680 e2.

44. Ziebuhr, J., The coronavirus replicase. Curr Top Microbiol Immunol, 2005. 287: p. 57–94.

45. Monto, A.S. and S.K. Lim, The Tecumseh study of respiratory illness. VI. Frequency of and relationship between outbreaks of coronavirus infection. J Infect Dis, 1974. 129(3): p. 271–6.

46. Shao, X., et al., Seroepidemiology of group I human coronaviruses in children. J Clin Virol, 2007. 40(3): p. 207–13.

47. Hofmann, H., et al., Human coronavirus NL63 employs the severe acute respiratory syndrome coronavirus receptor for cellular entry. Proc Natl Acad Sci U S A, 2005. 102(22): p. 7988–93.

48. Hamre, D. and M. Beem, Virologic studies of acute respiratory disease in young adults. V. Coronavirus 229E infections during six years of surveillance. Am J Epidemiol, 1972. 96(2): p. 94–106.

49. Chan, C.M., et al., Examination of seroprevalence of coronavirus HKU1 infection with S protein-based ELISA and neutralization assay against viral spike pseudotyped virus. J Clin Virol, 2009. 45(1): p. 54–60.

50. Dijkman, R., et al., The dominance of human coronavirus OC43 and NL63 infections in infants. J Clin Virol, 2012. 53(2): p. 135–9.

51. Dijkman, R., et al., Human coronavirus NL63 and 229E seroconversion in children. J Clin Microbiol, 2008. 46(7): p. 2368–73.

52. Kellam, P. and W. Barclay, The dynamics of humoral immune responses following SARS-CoV-2 infection and the potential for reinfection. J Gen Virol, 2020.

53. Vaisman-Mentesh, A., et al., SARS-CoV-2 specific memory B cells frequency in recovered patient remains stable while antibodies decay over time. medRxiv, 2020: p. 2020.08.23.20179796.

54. Sallusto, F., et al., From vaccines to memory and back. Immunity, 2010. 33(4): p. 451–63.

55. Kenney, L.L., et al., Increased Immune Response Variability during Simultaneous Viral Coinfection Leads to Unpredictability in CD8 T Cell Immunity and Pathogenesis. Journal of Virology, 2015. 89(21): p. 10786–10801.

56. Poh, C.M., et al., Two linear epitopes on the SARS-CoV-2 spike protein that elicit neutralising antibodies in COVID-19 patients. Nature Communications, 2020. 11(1): p. 2806.

57. Tylor, S., et al., The SR-rich motif in SARS-CoV nucleocapsid protein is important for virus replication. Can J Microbiol, 2009. 55(3): p. 254–60.

58. Surjit, M. and S.K. Lal, The SARS-CoV nucleocapsid protein: a protein with multifarious activities. Infect Genet Evol, 2008. 8(4): p. 397–405.

59. Li, F., Structure, Function, and Evolution of Coronavirus Spike Proteins. Annu Rev Virol, 2016. 3(1): p. 237–261.

60. Follis, K.E., J. York, and J.H. Nunberg, Furin cleavage of the SARS coronavirus spike glycoprotein enhances cell-cell fusion but does not affect virion entry. Virology, 2006. 350(2): p. 358–69.

61. Belouzard, S., et al., Mechanisms of coronavirus cell entry mediated by the viral spike protein. Viruses, 2012. 4(6): p. 1011–33.

62. Fung, T.S. and D.X. Liu, Human Coronavirus: Host-Pathogen Interaction. Annu Rev Microbiol, 2019. 73: p. 529–557.

63. Walls, A.C., et al., Unexpected Receptor Functional Mimicry Elucidates Activation of Coronavirus Fusion. Cell, 2019. 176(5): p. 1026–1039 e15.

64. Lai, A.L., et al., The SARS-CoV Fusion Peptide Forms an Extended Bipartite Fusion Platform that Perturbs Membrane Order in a Calcium-Dependent Manner. J Mol Biol, 2017. 429(24): p. 3875–3892.

65. Bradt, V., et al., Pre-existing yellow fever immunity impairs and modulates the antibody response to tick-borne encephalitis vaccination. npj Vaccines, 2019. 4(1): p. 38.

66. Andrews, S.F., et al., Immune history profoundly affects broadly protective B cell responses to influenza. Science Translational Medicine, 2015. 7(316): p. 316ra192–316ra192.

67. Diez, J.M., C. Romero, and R. Gajardo, Currently available intravenous immunoglobulin contains antibodies reacting against severe acute respiratory syndrome coronavirus 2 antigens. Immunotherapy, 2020. 12(8): p. 571–576.

68. Benner, S.E., et al., SARS-CoV-2 Antibody Avidity Responses in COVID-19 Patients and Convalescent Plasma Donors. J Infect Dis, 2020.

69. Bloch, E.M., et al., Deployment of convalescent plasma for the prevention and treatment of COVID-19. J Clin Invest, 2020. 130(6): p. 2757–2765.

70. Casadevall, A. and L.A. Pirofski, The convalescent sera option for containing COVID-19. J Clin Invest, 2020. 130(4): p. 1545–1548.

71. Zhou, G. and Q. Zhao, Perspectives on therapeutic neutralizing antibodies against the Novel Coronavirus SARS-CoV-2. Int J Biol Sci, 2020. 16(10): p. 1718–1723.

72. Zhang, Q., et al., Inborn errors of type I IFN immunity in patients with life-threatening COVID-19. Science, 2020: p. eabd4570.

73. Madeira, F., et al., The EMBL-EBI search and sequence analysis tools APIs in 2019. Nucleic Acids Res, 2019. 47(W1): p. W636–W641.

74. Katoh, K. and D.M. Standley, MAFFT multiple sequence alignment software version 7: improvements in performance and usability. Mol Biol Evol, 2013. 30(4): p. 772–80.

75. Madeira, F., et al., Using EMBL-EBI Services via Web Interface and Programmatically via Web Services. Curr Protoc Bioinformatics, 2019. 66(1): p. e74.

76. Dhanda, S.K., et al., IEDB-AR: immune epitope database-analysis resource in 2019. Nucleic Acids Res, 2019. 47(W1): p. W502–W506.

