## Supplementary Material for "Distinct antibody repertoires against endemic human coronaviruses in children and adults"

Version 2, 20 September 2020

#### Supplementary Figures

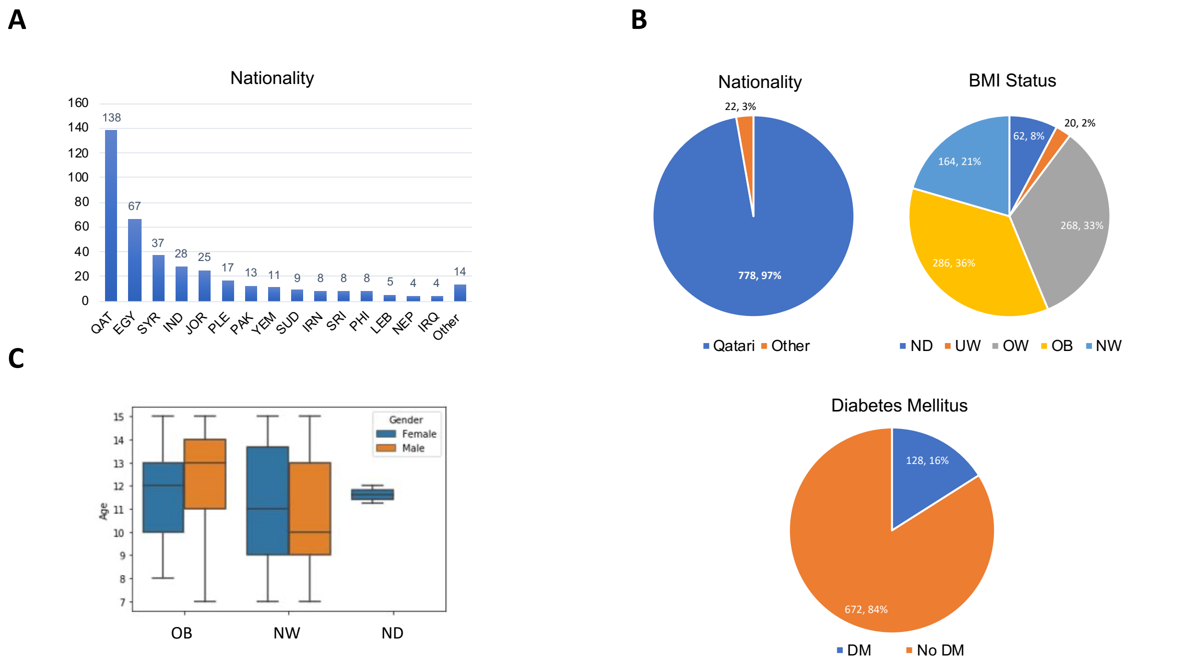


**Supplementary Figure S1. Demographic and clinical features of the different human cohorts. A,** Nationalities among the healthy blood bank donors. **B,** Number and fraction of Qatari nationals and long-term residents with other nationality comprising the 800 assayed individuals of the Qatar Biobank cohort, stratified by body mass index (BMI) and proportion of subjects with diabetes mellitus (DM). UW, underweight (BMI < 18); NW, normal weight (BMI ≥ 18 and < 25); OW, overweight (BMI ≥ 25 and <30); OB, obese (BMI ≥ 30). **C,** Number and fraction of male and female pediatric study subjects with normal weight (NW) or who were obese (OB). ND, not determined; QAR, Qatar; EGY, Egypt; SYR, Syria; IND, India; JOR, Jordan; PAK, Pakistan; YEM, Yemen; SUD, Sudan; IRN, Iran; SRI, Sri Lanka; PHI, Philippines; LEB, Lebanon; NEP, Napal; IRQ, Iraq; Other, residents with other nationality; PLE, non-residents.

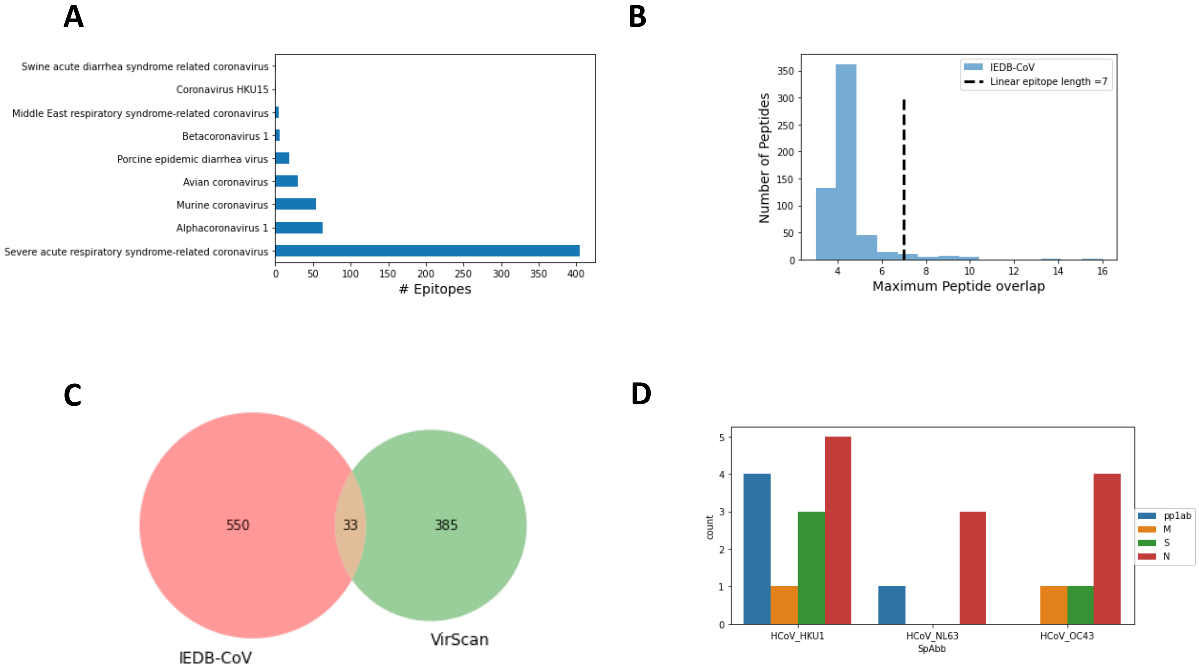


**Supplementary Figure S2. Comparison analysis of enriched HCoV peptides identified by PhIP-Seq with B cell epitopes in IEBD ([www.iebd.org](www.iebd.org)). A**, Number of B cell epitopes for different CoV species available in IEBD (accessed and downloaded from www.iedb.org on 17 May 2020). **B,** Number of peptides from endemic HCoVs which we found to be significantly enriched in at least three of all 1399 samples with homology in the linear amino acid sequence to 583 known CoV B cell epitopes in IEBD. We considered an epitope as a match if there was ≥ 7 linear sequence identity between query (IEBD epitope) and the amino acid sequence of the enriched HCoV peptide. **C,** Venn diagram depicting overlap between enriched peptides (i.e. matches) detected by VirScan and B cell epitopes from CoVs in IEDB. **D,** Number of matching peptides that share sequence homology with known B cell epitopes, stratified by species and viral protein.

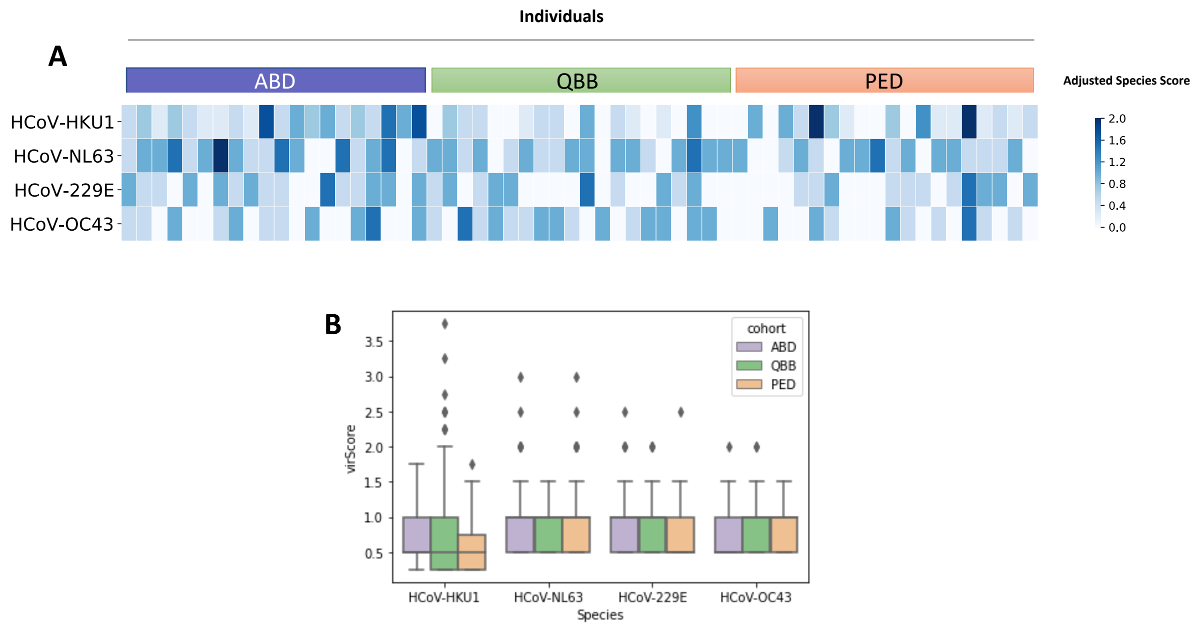


**Supplementary Figure S3: Similar antibody repertoire breadth among seropositive individuals of the different cohorts. A,** Heatmap plot depicting adjusted species score values for HCoV-HKU1, -NL63, -229E and -OC43 in randomly selected seropositive individuals (n = 20) of each cohort. Each individual is represented as a column. Values ≥ 1 indicate that the individual was seropositive for a given species. **B,** Species-wise variation in adjusted species score values among seropositive individuals of each cohort.

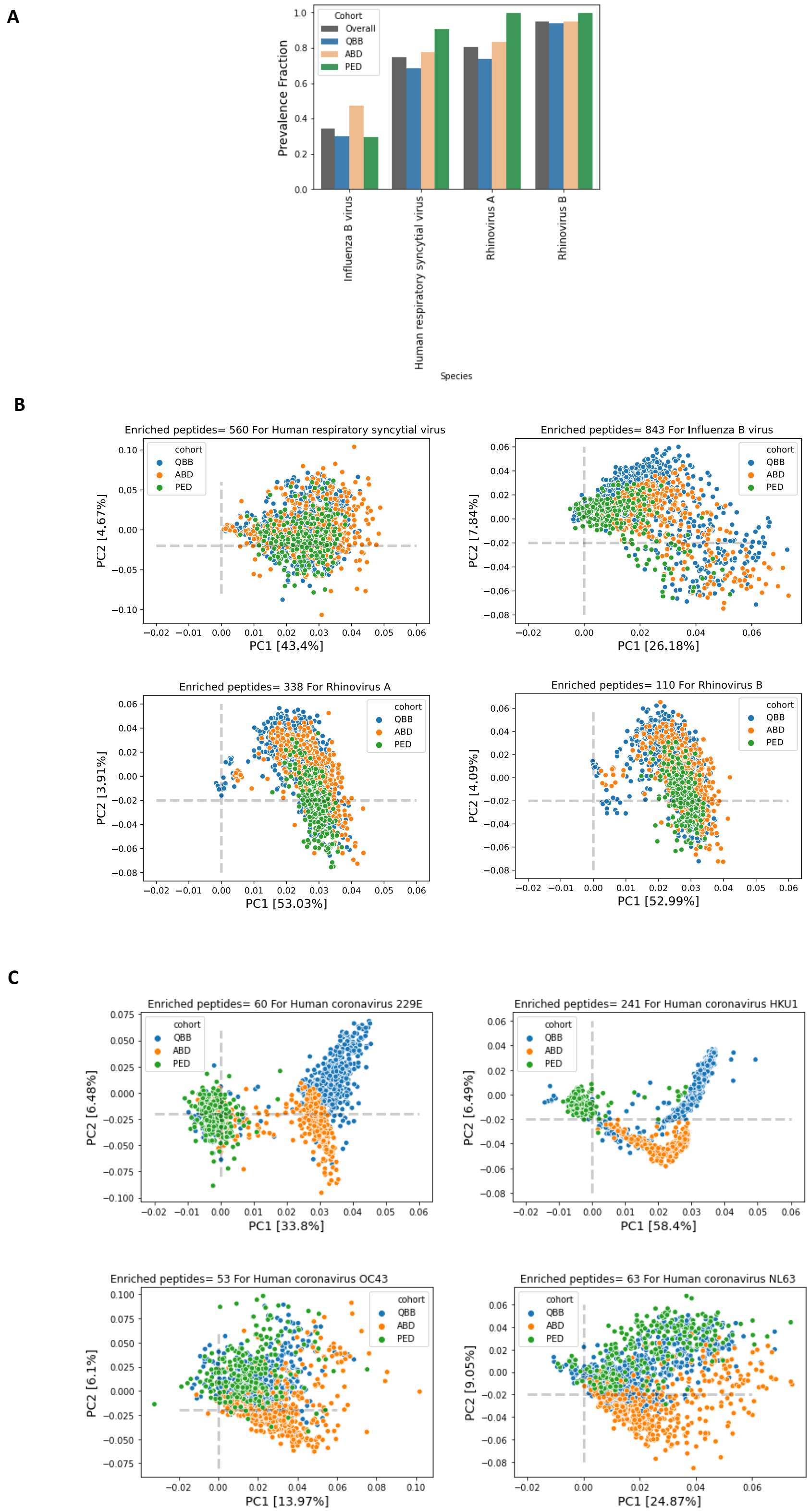


**Supplementary Figure S4. Seroprevalence of respiratory viruses and variance in the antibody repertoires among individuals of the analyzed cohorts. A,** Bar plot depicting the fraction of individuals found seropositive for human respiratory syncytial virus, rhinovirus A, rhinovirus B and influenza B virus among the tested subjects or by cohort. **B and C,** Principal component analysis of the antigenic peptides of human respiratory syncytial virus, rhinovirus A, rhinovirus B and influenza B **(B)** as well as of human coronaviruses 229E, NL63, OC43 and HKU1 **(C)** that we found to be enriched in at least 3 samples.

A

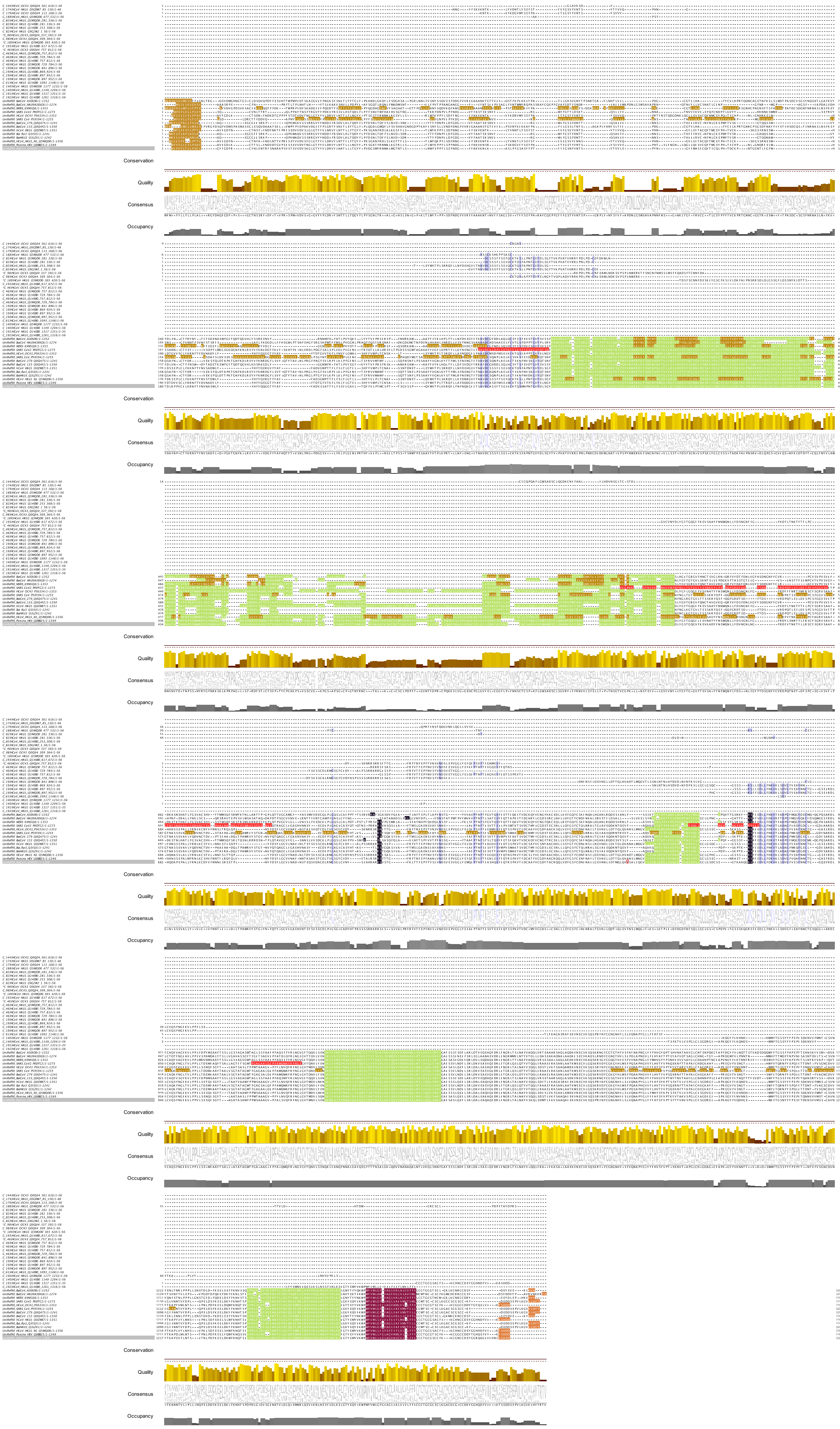

B

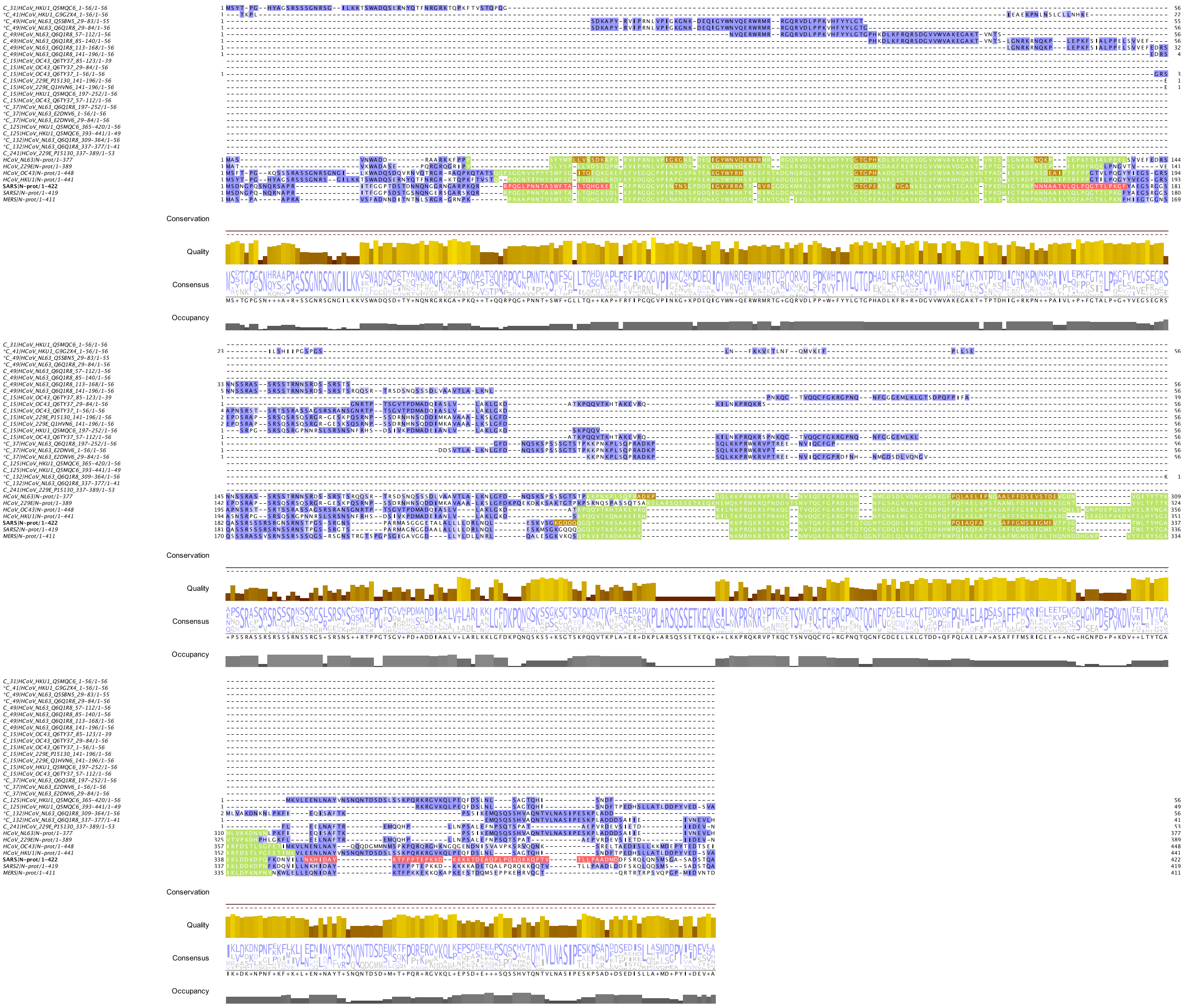

**C**

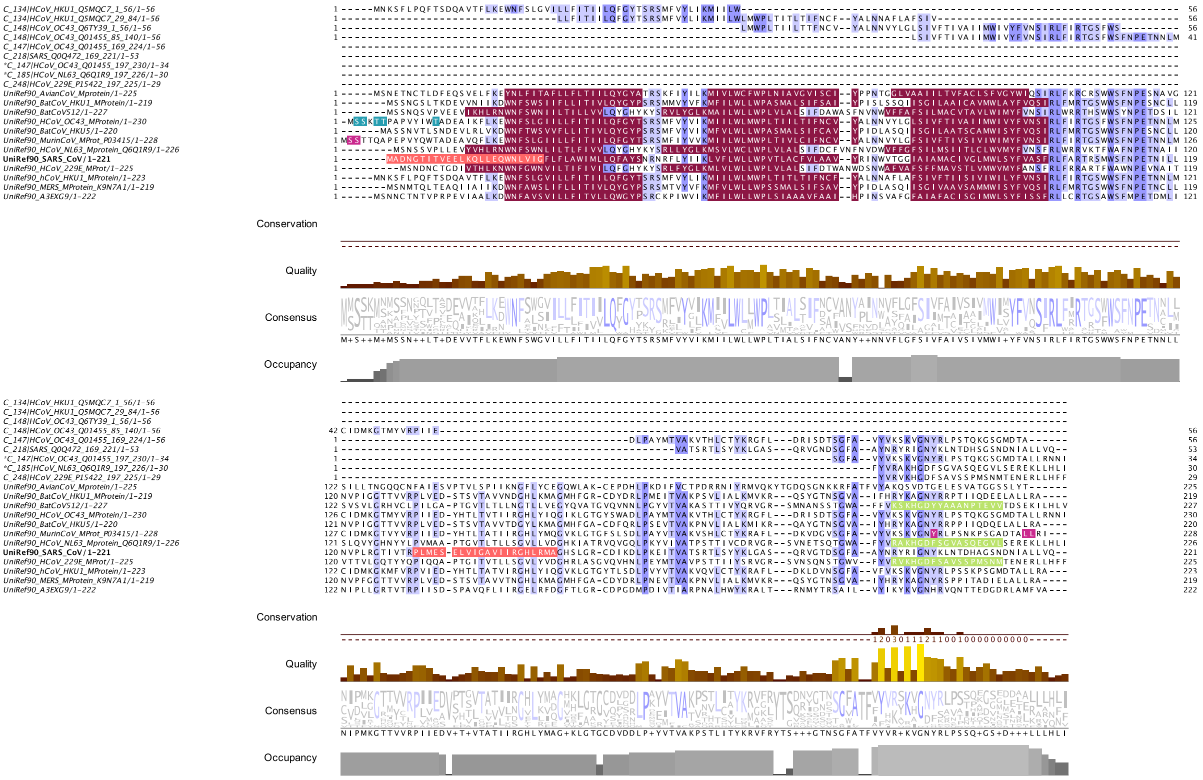
Conservation
Quality
Consensus
Occupancy
UniRef90_SARS_CoV/1-221

**D**

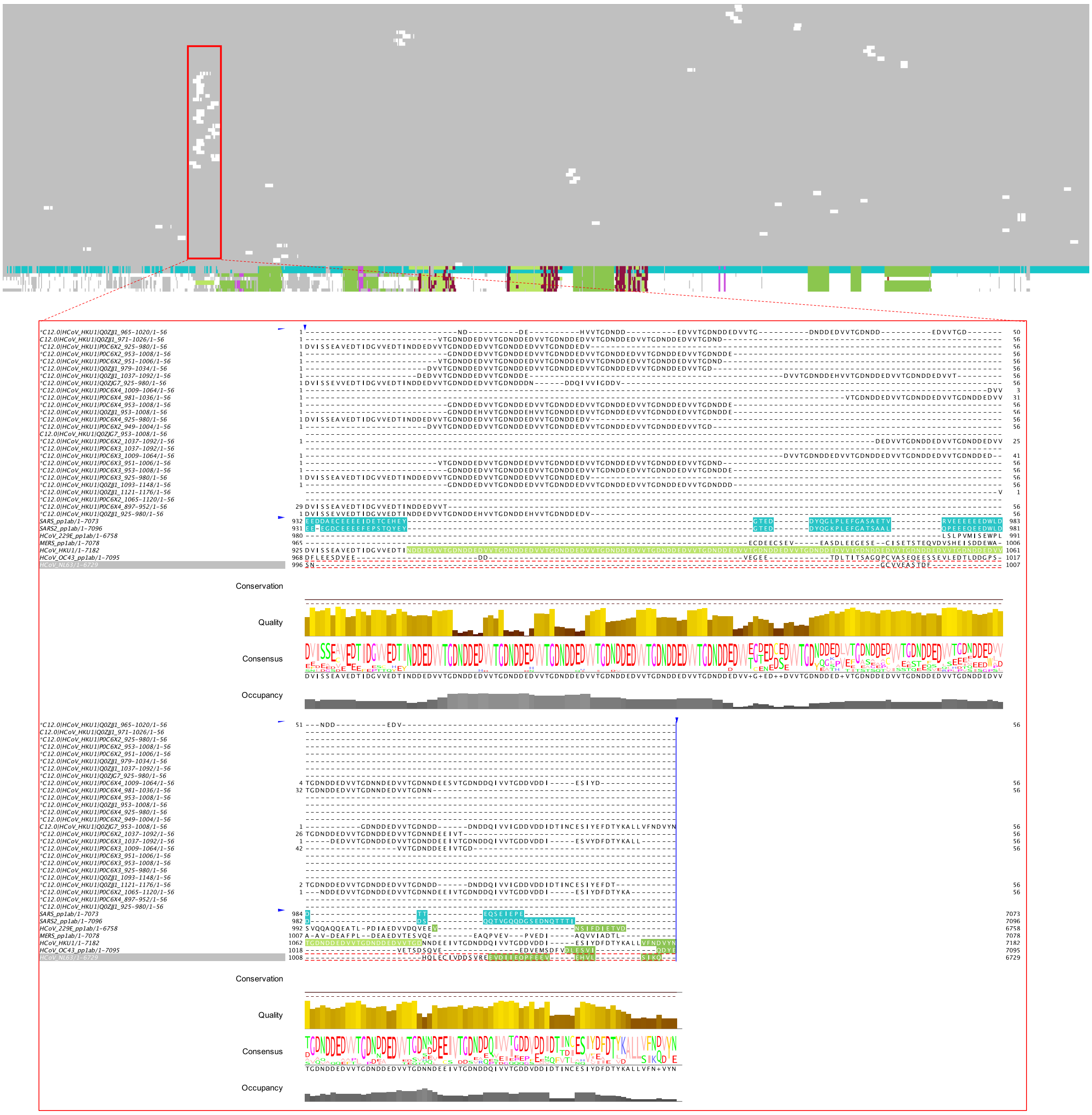

**Supplementary Figure S5. Mapping of enriched, immunodominant peptides of endemic HCoVs to the full-length protein sequences**. **A-D**, Multiple sequence alignments of the enriched, immunodominant peptides with the full-length sequences of the spike protein (S), nucleocapsid protein (N), matrix glycoprotein (M) and the Orf1ab replicase polyprotein (pp1ab) of selected CoVs. Amino acids with high percentage of sequence identity are shown in violet. Reference sequence features are color-coded in accordance to the annotation and features available in UniProt (subjected to availability) and visualized using JalView. Green indicates regions of interest, red indicates transmembrane regions, orange indicates beta strands and pink indicates residues annotated as epitopes in IEDB. The cleavage site (S1/S2 and S2) of the SARS-CoV S protein is shaded in black. For pp1ab, the sequence alignment is shown as an overview with gaps are shaded in grey and sequences are in white. Only part of the sequence alignment is shown, which depicts the PL1-PRO cleavage product containing a tandem repeat of N-[DN]-D-E-D-V-V-T-G-DA in HCoV-HKU1 (isolate N1) (UniProt entry and position P0C6X2[945-1084]).

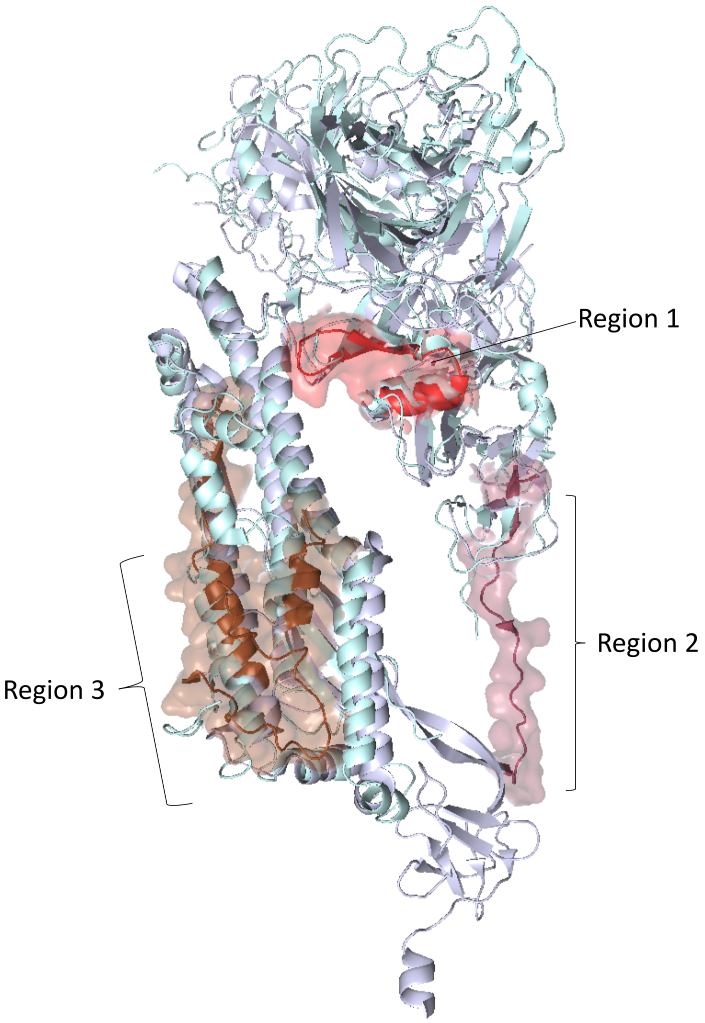


**Supplementary Figure S6. Superimposition of Spike protein of SARS-CoV2 (6VXX) and HCoV-HKU1 (5I08) in the prefusion state.** Root-mean-square deviation [rmsd] = 3.328 Ångström.

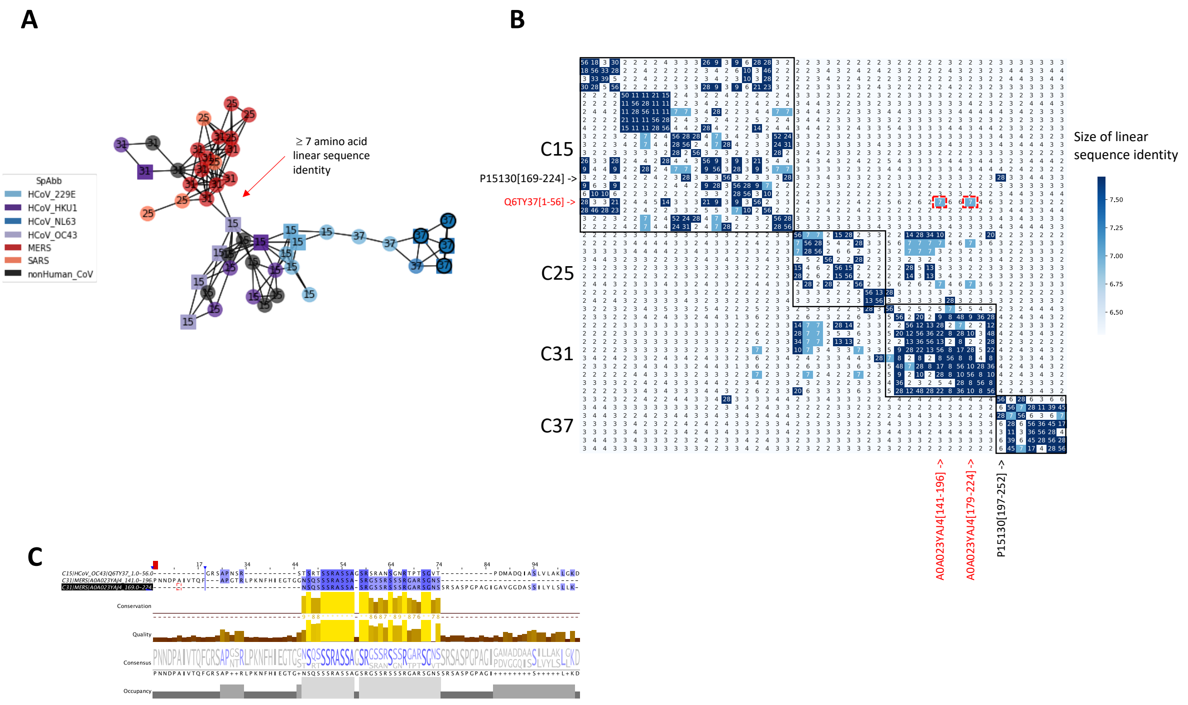


**Supplementary Figure S7. Clustering of peptides for shared linear B cell epitopes. A,** Network representation of peptides built from a pairwise distance matrix as shown in Figure 5B. Each node represents an enriched peptide and the color indicates the species. Edges indicate ≥ 7 amino acids linear sequence identity between two nodes (i.e. peptides), the estimated size of a linear B cell epitope. **B,** Pairwise distance matrix of peptides targeted by cross-reactive antibodies, including enriched peptides of clusters 15, 25 ,31 and 37. The cluster assignment has been indicated as a black box. Distance pairs with ≥ 7 amino acid linear sequence identity are shown in dark blue. Distance pairs marked with a red box represent a 7 amino acid linear sequence identity between enriched peptides of more distantly related CoVs, namely HCoV-OC43 (Q6TY37[1-56]) and MERS-CoV (A0A023YAJ4[141-196] and A0A023YAJ4[179-224]). **C,** Multiple sequence alignment of enriched HCoV-OC43- and MERS-CoV-derived peptides Q6TY37[1-56], A0A023YAJ4[141-196] and A0A023YAJ4[179-224] encoding part of the N-terminal intrinsically disordered region (IDR) of the N protein. The alignment shows the 7 amino acid linear sequence identity in a low-complexity sequence motive (SSRASSA) which likely represents a broadly cross-reactive antibody binding site.

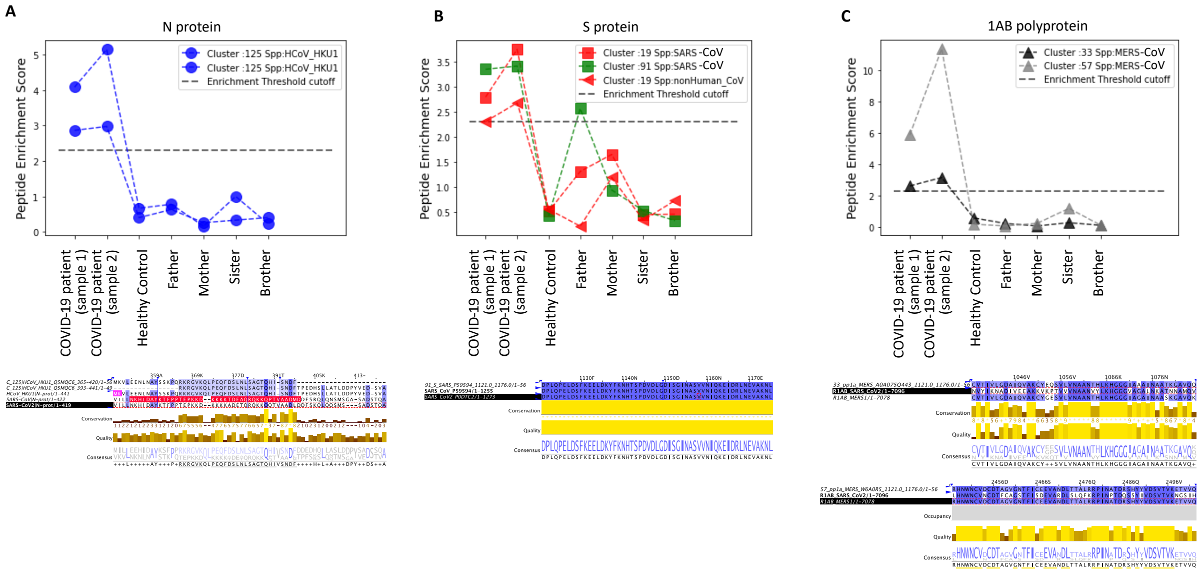


**Supplementary Figure S8. Cross-reactive antibody responses in a COVID-19 patient with proteins of other coronaviruses.** Plasma samples of an adult COVID-19 patient, unexposed family members (father, mother, sister and brother) as well as an unrelated healthy, age- and gender-matched control were subjected to PhIP-Seq analysis. Samples from the COVID-19 patient were obtained at two time points: during acute infection (sample 1) and 30 days after infection (sample 2). Shown is a comparison of enrichment scores of peptides derived from the N protein (A), S protein (B) and 1AB polyprotein (C) of various coronaviruses. HCoV-HKU1 peptides are depicted as circles, SARS-CoV peptides are depicted as squared symbols and MERS-CoV as well as non-human CoV peptides are shown as triangle shapes. SARS-CoV-2 peptides are not included in the VirScan phage library used in our study. The dashed line indicates the significance cut-off for the peptide enrichment [−log10(P) ≥ 2.3]. The cluster numbers indicate linear epitopes as listed in Table S3. The multiple sequence alignments (bottom) of the enriched peptides with the reference sequences of SARS-CoV-2 and the respective CoV species show the degree of sequence identity across species. Known epitopes (IEBD) are shown in pink color. Sequences are colored to indicate percent sequence identity using Jalview.

### Supplementary Tables

**Supplementary Table S1.** Proteins of CoVs represented in the VirScan phage library. Endemic hCoVs are shown in bold.

| *Species* | *Organism* | *Protein* | *Number of Peptides* |
|---|---|---|---|
| *Alphacoronavirus 1* | Feline coronavirus (strain FIPV WSU-79/1146) (FCoV) | NS | 239 |
| *Bat coronavirus 1B* | Bat coronavirus 1B | Rep1a | 249 |
| *Betacoronavirus 1* | Bovine coronavirus | N | 4 |
| | Bovine coronavirus (strain 98TXSF-110-ENT) (BCoV-ENT) (BCV) | E | 2 |
| | | N | 7 |
| | | NS | 12 |
| | | S | 48 |
| | Equine coronavirus | Rep1a | 254 |
| | **Human coronavirus OC43 (HCoV-OC43)** | **HE** | **15** |
| | | **M** | **11** |
| | | **N** | **4** |
| | | **NS** | **253** |
| | | **S** | **37** |
| | Porcine hemagglutinating encephalomyelitis virus (strain 67N) (HEV-67N) | N | 16 |
| ***Human coronavirus 229E*** | **Human coronavirus 229E (HCoV-229E)** | **E** | **2** |
| | | **M** | **8** |
| | | **N** | **23** |
| | | **NS** | **248** |
| | | **ORF4** | **10** |
| | | **Rep1b** | **2** |
| | | **S** | **63** |
| ***Human coronavirus HKU1*** | **Human coronavirus HKU1 (HCoV-HKU1)** | **N** | **13** |
| | | **Rep1ab** | **510** |
| | | **S** | **15** |
| | **Human coronavirus HKU1 (isolate N1) (HCoV-HKU1)** | **E** | **2** |
| | | **HE** | **13** |
| | | **M** | **7** |
| | | **N** | **22** |
| | | **NS** | **259** |
| | | **S** | **48** |
| | **Human coronavirus HKU1 (isolate N2) (HCoV-HKU1)** | **E** | **2** |
| | | **HE** | **13** |
| | | **N** | **7** |
| | | **NS** | **258** |
| | | **S** | **48** |
| | **Human coronavirus HKU1 (isolate N5) (HCoV-HKU1)** | **NS** | **254** |
| ***Human coronavirus NL63*** | **Human coronavirus NL63 (HCoV-NL63)** | **E** | **2** |
| | | **M** | **10** |
| | | **N** | **18** |
| | | **NS** | **248** |
| | | **ORF3** | **1** |
| | | **Rep1a** | **1** |
| | | **S** | **71** |
| ***Middle East respiratory syndrome coronavirus*** | BtVs-BetaCoV/SC2013 | E | 2 |
| | | M | 7 |
| | | N | 15 |
| | | ORF3 | 3 |
| | | ORF4 | 11 |
| | | ORF5 | 8 |
| | | ORF8 | 6 |
| | | Rep1a | 256 |
| | | S | 47 |
| | Coronavirus Neoromicia/PML-PHE1/RSA/2011 | E | 2 |
| | | ORF3 | 3 |
| | | ORF4 | 12 |
| | | ORF5 | 7 |
| | | ORF8 | 7 |
| | | Rep1a | 156 |
| | | S | 47 |

| | | | |
|---|---|---|---|
| | Human coronavirus EMC (isolate United Kingdom/H123990006/2012) (HCoV-EMC) | E | 2 |
| | | M | 7 |
| | | N | 14 |
| | | NS | 408 |
| | | ORF3 | 3 |
| | | ORF4 | 11 |
| | | ORF5 | 7 |
| | | S | 48 |
| | Middle East respiratory syndrome coronavirus | N | 8 |
| | | ORF4 | 11 |
| | | ORF5 | 11 |
| | | ORF8 | 3 |
| | | Rep1a | 1018 |
| | | Rep1ab | 95 |
| | | Rep1b | 63 |
| | | S | 108 |
| *Severe acute respiratory syndrome-related coronavirus* | Bat coronavirus 279/2005 (BtCoV) (BtCoV/279/2005) | E | 2 |
| | | M | 7 |
| | | NS | 254 |
| | | U | 2 |
| | Human SARS coronavirus (SARS-CoV) (Severe acute respiratory syndrome coronavirus) | N | 15 |
| | | NS | 9 |
| | | ORF3 | 9 |
| | | ORF7 | 4 |
| | | ORF9 | 3 |
| | | S | 44 |
| | Human SARS coronavirus (isolate Tor2) (SARS-CoV) (Severe acute respiratory syndrome coronavirus) | Rep1b | 1 |

**Supplementary Table S2.** Number of individuals per cohort included in the downstream analysis.

| Cohort | Gender | Number | Age (years) | | | |
|---|---|---|---|---|---|---|
| | | | Mean | Std | Min | Max |
| ABD | Male | 370 | 40.1 | 10.1 | 19 | 66 |
| PED | Female | 117 | 11.4 | 2.2 | 7 | 15 |
| | Male | 114 | 11.8 | 2.3 | 7 | 15 |
| QBB | Female | 511 | 40.4 | 12.9 | 19 | 81 |
| | Male | 287 | 41.0 | 12.9 | 19 | 81 |

QBB, Qatar Biobank cohort; ABD, adult blood bank donors; PED, pediatric subjects.

**Supplementary Tables S3 and S4. List of enriched HCoV peptides and cluster assignment.**

<TableS3 and S4_2020-8-27.xls>
